# A snapshot of the Physcomitrella N-terminome reveals N-terminal methylation of organellar proteins

**DOI:** 10.1101/2024.06.11.598444

**Authors:** Sebastian N.W. Hoernstein, Andreas Schlosser, Kathrin Fiedler, Nico van Gessel, Gabor L. Igloi, Daniel Lang, Ralf Reski

## Abstract

Post- or co-translational N-terminal modifications of proteins influence their half-life as well as mediating protein sorting to organelles via cleavable N-terminal sequences that are recognized by the respective translocation machinery. Here, we provide an overview on the current modification state of the N-termini of over 4500 proteins from the model moss Physcomitrella (Physcomitrium patens) using a compilation of 24 N-terminomics datasets. Our data reveal distinct proteoforms and modification states and confirm predicted targeting peptide cleavage sites of 1144 proteins localized to plastids and the thylakoid lumen, to mitochondria, and to the secretory pathway. Additionally, we uncover extended N-terminal methylation of mitochondrial proteins. Moreover, we identified PpNTM1 (P. patens alpha N-terminal protein methyltransferase 1) as a candidate for protein methylation in plastids, mitochondria and the cytosol. These data can now be used to optimize computational targeting predictors, for customized protein fusions and their targeted localization in biotechnology, and offer novel insights into potential dual targeting of proteins.

**Key message:** Analysis of the N-terminome of Physcomitrella reveals N-terminal monomethylation of nuclear encoded, mitochondria-localized proteins.

**Graphical abstract:** 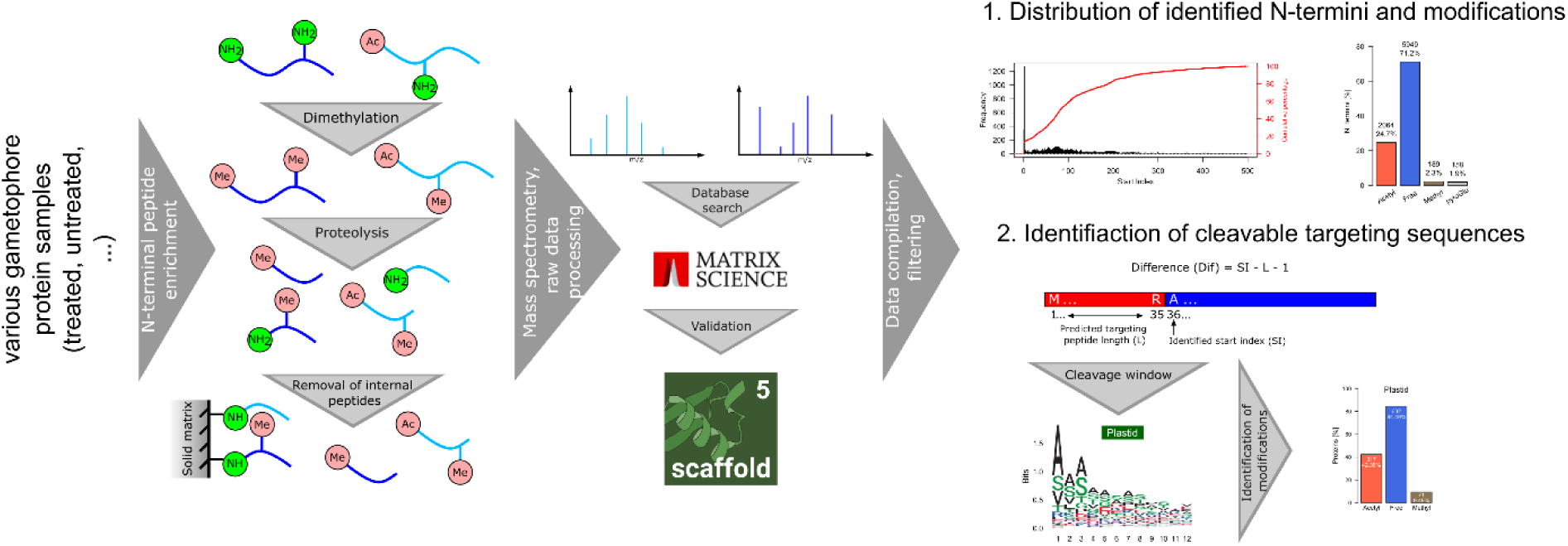

## Introduction

Following translation at the ribosome, the N-terminus of a protein is subjected to a plethora of modifications among which are proteolytic processing and the addition of moieties such as acetyl, methyl or other functional groups (Meinnel and Giglione 2008; Fortelny et al. 2015). In turn, the N-terminus enables subcellular targeting and determines protein half-life (Varshavsky 1996; Kunze and Berger 2015; Armenteros et al. 2019; Varshavsky 2019). Modifications of the N-terminus are introduced in a co- or a posttranslational manner with cotranslational acetylation and methionine-excision being among the most abundant modifications in eukaryotes (Ree et al. 2018; Giglione and Meinnel 2021). In plants, however, N-terminal acetylation also occurs in a posttranslational manner on plastid stromal proteins after import and cleavage of their targeting peptide (Giglione and Meinnel 2021). In contrast to proteolytic trimming, amino acids can also be added to the apparent N-terminus of a protein in a ribosome-independent manner (Varshavsky 1996; Tasaki et al. 2012) as part of the N-degron pathway for targeted proteolysis. Various methods such as COFRADIC (Staes et al. 2011), TAILS (Kleifeld et al. 2010) or HUNTER (Demir et al. 2022) have been established and permit the characterization of proteases, high-throughput degradomics and profiling of N-terminal acetylation. In turn, N-terminomics data are available from public databases such as TopFIND (https://topfind.clip.msl.ubc.ca) for various organisms including human, mouse, yeast, and Arabidopsis.

In contrast, almost no N-terminomics data were available for the model plant Physcomitrella (*Physcomitrium patens*; Lueth and Reski 2023). This moss is a versatile model system for evo-devo studies (Horst et al. 2016), plant physiology (Decker et al. 2017; Wiedemann et al. 2018) and evolution of metabolic pathways (Renault et al. 2017; Knosp et al. 2024) due to its interesting evolutionary position at the early divergence of land plants (Rensing et al. 2008). It has further proven to be a valuable system for proteomic and proteogenomic research due to its easy and axenic culture conditions (Hohe et al. 2002) enabling highly reproducible and even GMP-compliant culture conditions (Sarnighausen et al. 2004; Heintz et al. 2006; Mueller et al. 2014; Hoernstein et al. 2016, 2018; Fesenko et al. 2019, 2021). Besides broad application in basic research, Physcomitrella is employed as a production platform for recombinant biopharmaceuticals in GMP-compliant bioreactors (Decker and Reski 2020; Ruiz-Molina et al. 2022; Tschongov et al. 2024). Furthermore, genomic and transcriptomic resources are well established and publicly available (Lang et al. 2018; Perroud et al. 2018; Fernandez-Pozo et al. 2020; Bi et al. 2024).

Here, we provide a snapshot of the N-terminome of the moss Physcomitrella with a focus on the cleavage of N-terminal targeting sequences, N-terminal acetylation and N-terminal monomethylation. The data was compiled using 24 datasets from various experimental setups and subsequent N-terminal peptide enrichment using a modified TAILS approach. We reveal apparent N-terminal methylation not only of plastid and cytosolic proteins but also of mitochondrial proteins. Furthermore, we provide a list of confirmed targeting peptide cleavage sites along with a candidate list of proteins which are dually targeted to plastids and mitochondria as well as to mitochondria and the cytosol. With this, we provide a resource for basic research as it contains information about translation of splice variants as well as posttranslational and posttranscriptional processing of proteins. Moreover, targeting of recombinant proteins to plastids of *Nicotiana benthamiana* (Maclean et al. 2007) or the extracellular space in Physcomitrella (Schaaf et al. 2005) enabled high yields of the desired recombinant product. Consequently, our present data also provide a comprehensive resource for further customized recombinant protein production and targeting in Physcomitrella.

## Results and Discussion

### Overview of identified N-termini

The present data provide a qualitative overview of the N-terminome of the moss Physcomitrella using a compilation of 24 datasets from N-terminal peptide enrichments from different tissues, treatments and different sample processing protocols. The datasets were obtained during method establishment for various purposes not related to the analysis performed in this study and hence the data was only assessed qualitatively and will not allow any cross-sample comparison. A table providing details about the sample type, tissue employed and other experimental parameters is available from Supplemental Table S1.

Enrichment of N-terminal peptides was performed as described in Hoernstein et al. (2018) with modifications. Free amino groups in the protein sample were blocked by reductive dimethylation according to Kleifeld et al. (2010) and depletion of internal peptides after proteolysis was performed according to McDonald and Beynon (2006). Mass spectrometry (MS) measurements were performed on an LTQ-Orbitrap Velos Pro (ThermoScientific, Waltham, MA, USA) and raw data were processed and searched with *Mascot* (Matrix Science, Chicago, IL, USA). All database search results were loaded in *Scaffold5^TM^* (V5.0.1, https://www.proteomesoftware.com) software and proteins were accepted with a *ProteinProphet^TM^* (Nesvizhskii et al. 2003) probability of at least 99% and a minimum of 1 identified peptide. Peptides were accepted at a *PeptideProphet^TM^* (Keller et al. 2002) probability of at least 95% and a *Mascot* ion score of a least 40 (Supplemental Table S2). Using these settings, from a total of 24 datasets we identified 11,533 protein N-termini using 32,213 spectra corresponding to 4517 proteins (3920 protein groups) with a decoy FDR (false discovery rate) of 0.4% at the protein level and 0.08% at the peptide level (Supplemental Tables S2, S3). Approximately 20% of the identified N-termini represented either the initiator methionine (start index 1, Figure 1A) or the subsequent amino acid after cleavage of the initiator methionine (start index 2). For approximately 40% a start index between 2 and 100 was identified, indicating proteolytic processing and cleavage of subcellular targeting sequences. A single experimentally determined N-terminus was observed in approximately 70% of all cases whereas for approximately 30% of the identified proteins two or more N-termini were observed (Figure 1B). This is in strong contrast to previous findings from proteins in Physcomitrella bioreactor supernatants (Hoernstein et al. 2018) where at least two distinct N-termini were observed for approximately 80% of all identified proteins. Further, we analyzed the presence of N-terminal modifications with a focus on N-terminal acetylation, monomethylation and presence of pyro-glutamate (pyroGlu) at the N-terminus. The latter modification can occur spontaneously or *via* enzymatic catalysis on N-terminal glutamine residues (Schilling et al. 2008). Since pyro-glutamate formation can also occur following proteolysis during sample processing (Purwaha et al. 2014), only peptides where the preceding P1 amino acid did not match the specificity of the employed protease (e.g., no peptides with K or R as preceding amino acid in the case of trypsin digests) were considered here.

**Figure 1.**
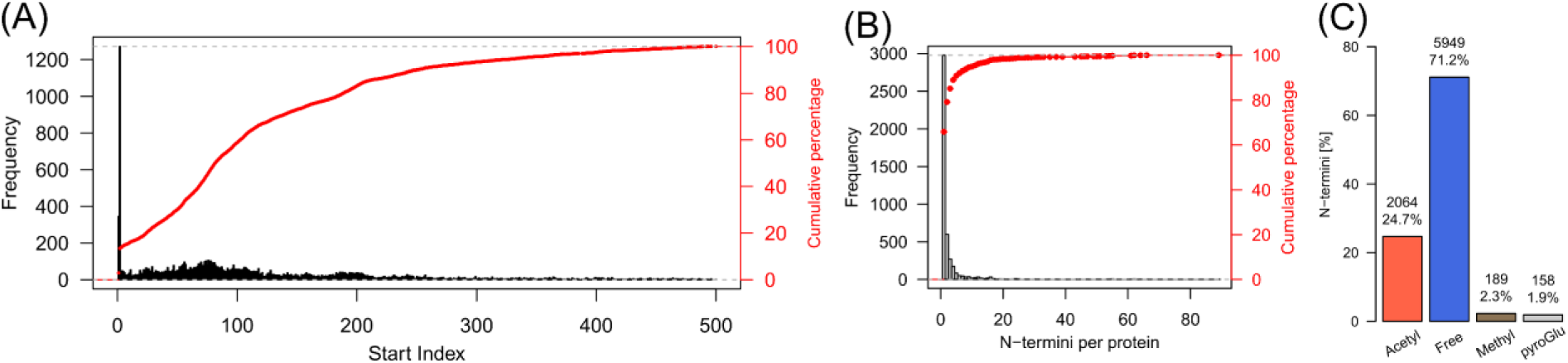
Overview of identified N-termini, N-terminal modifications and identified cleavage sites of targeting peptides. (**A**) Frequency of identified N-terminal positions per identified protein accession. The start index represents the position number of the identified N-terminal amino acid in the corresponding protein model. (**B**) Frequency of the number of identified N-termini per protein. (**C**) Bar chart depicting the distribution of identified N-terminal modifications. Peptides bearing N-terminal pyro-glutamate (pyroGlu) were only counted if the preceding amino acid did not match the specificity of the applied protease (e.g., peptides identified with K|R in the P1 position were rejected in the case of trypsin digests).

Approximately 25% of all identified N-termini were acetylated, 71% had no modification and 4% were either methylated or had N-terminal pyroGlu (Figure 1C). The actual level of N-terminal pyroGlu occurrence is likely higher, but due to specificity ambiguity with the experimentally employed proteases this cannot be analyzed further. At the protein level, we found approximately 76% of the nuclear-encoded proteins having either the retained or the cleaved initiator methionine (1436 protein groups, Supplemental Table S3) to be N-terminally acetylated (1097 protein groups, Supplemental Table S3). This degree is slightly below the estimated degree of around 90% of N-terminal acetylated proteins in plants (Bienvenut et al. 2012; Linster and Wirtz 2018).

### Post-import trimming of plastid proteins

Cleavable N-terminal sequences are required for sub- and extracellular targeting of nuclear-encoded proteins. Their cleavage *via* specific proteases after translocation across a respective organellar membrane generates a new N-terminus that represents either the final N-terminus of the translocated protein or a new site for further proteolytic processing by organellar proteases. For Physcomitrella, a total of 8681 cleavable N-terminal targeting sequences are predicted (Supplemental Figure S1) and here we compared our experimentally observed N-termini to these predictions allowing a tolerance window of ±5 amino acids around a predicted targeting peptide cleavage site (Figure 2A). In the following, a difference of 0 indicates agreement of an observed N-terminal amino acid with a predicted cleavage site (predicted P1 amino acid, Figure S2A). Within this range we confirm the predicted cleavage sites of 748 plastid targeting signals (cTP), of 57 thylakoid luminal targeting signals (luTP), of 154 mitochondrial presequences (mTP), and of 185 secretory signal peptides (SP) using our present N-terminomics data (Figure S2B). This data is compiled in Supplemental Table S4.

**Figure 2.**
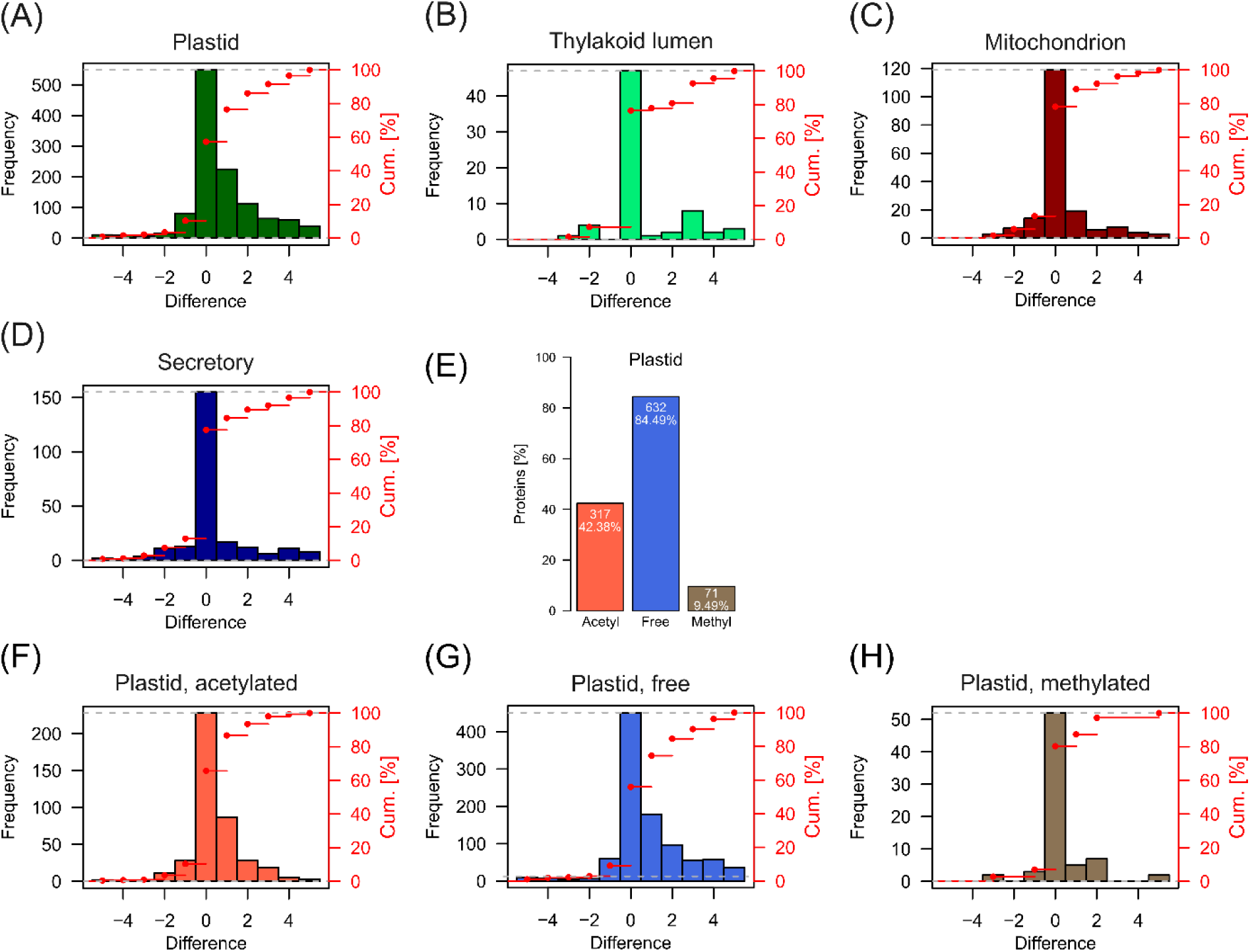
Comparison of experimentally observed N-termini with predicted organellar targeting peptide cleavage sites. Depicted are frequencies of identified N-termini around a predicted targeting peptide cleavage site. A difference of 0 indicates an identified N-terminal amino acid corresponds to the P1’ amino acid of a predicted cleavage site. Cleavages of plastid (**A**), thylakoid lumen (**B**), mitochondrial (**C**) and secretory (**D**) targeting signals were predicted with *TargetP2.0*. All data are available from Supplemental Table S4. (**E**) Bar chart depicting the distribution of plastid protein isoforms with confirmed cleavage of a plastid targeting peptide and their identified N-terminal modifications. Percentages are related to the total number of identified proteins with a cleaved N-terminal plastid targeting peptide (748) within a window of ±5 amino acids around a predicted cleavage site. All data are available from Supplemental Table S4. Frequency of identified N-termini around a predicted plastid transit peptide cleavage site being either acetylated (**F**), unmodified (free, (**G**)) or monomethylated (**H**). Cum. [%]: cumulative percentage (red points).

Among the confirmed plastid proteins, we find approximately 42% to be N-terminally acetylated (317 protein isoforms, Supplemental Table S4). Apparently, N-termini identified around a plastid transit peptide cleavage site only matched the predicted cleavage site exactly for approximately 47% (Figure 2A). This percentage is strikingly higher in all other cases with almost 70% for thylakoid luminal transit peptides (Figure 2B) and around 65% for mitochondrial presequences and secretory signal peptides (Figures 2C, D). Further, approximately 43% of the N-termini of plastid proteins within the chosen difference window deviate by one to five amino acids upstream of the predicted cleavage site with decreasing frequency. This effect is less apparent for luminal targeting sequences (approximately 33%), mitochondrial presequences (approximately 22%) and secretory signal peptides (approximately 32%). Consequently, the distribution of differences between predicted and observed plastid transit cleavage site (Figure 2A) indicates a successive proteolytic postprocessing pattern of plastid proteins after cleavage of their transit peptide. A similar scenario with multiple cleavage sites around predicted plastid transit peptide cleavage sites has also been observed in Arabidopsis (Bienvenut et al. 2012; Rowland et al. 2015).

### N-terminal modifications of plastid and mitochondrial proteins

Among the proteins with an identified plastid transit peptide cleavage site, we found 42% protein isoforms with an acetylated N-terminus and almost 10% with a monomethylated N-terminus (Figure 2E). Strikingly, plastid N-termini being acetylated and non-modified (free) both show this successive cleavage pattern whereas monomethylated N-termini do not show this pattern (Figures 2F-H).

This raises the question whether this processing occurs on both, acetylated and free N-termini, or whether only free N-termini are processed and subsequently acetylated. One explanation would be that N^α^-acetylation of plastid proteins is incomplete and affects only a fraction of each protein isoform. Effectively, many N-termini of plastid proteins were identified in this study in a dual state, being acetylated and free (e.g., Pp3c18_19140V3.1, Pp3c15_7750V3.4; Supplemental Table S4). In this case, N^α^-acetylation would prevent N-terminal trimming, whereas the fraction with an unmodified N-terminus would be proteolytically processed to different levels and subsequently acetylated. This scenario may be supported by the fact that N^α^-acetylated and free N-termini share a similar preference of N-terminal amino acids (Figure 3), with alanine and serine being the most prominent ones. Apparently, the relative amino acid frequency of the N-terminally acetylated plastid proteins is strikingly similar to the relative frequency in Arabidopsis (Huesgen et al. 2013). Although the plastid protease inventory is under active investigation (Meinnel and Giglione 2022; van Wijk 2024), a specific protease for such N-terminal trimming has not yet been identified. Aminopeptidases identified in Arabidopsis were recently proposed to also confer trimming functions (Rowland et al. 2015; Meinnel and Giglione 2022), but their activity was investigated on released plastid transit peptides in conjunction with presequence proteases (Teixeira et al. 2017), and not on the protein N-terminus of the corresponding protein.

**Figure 3.**
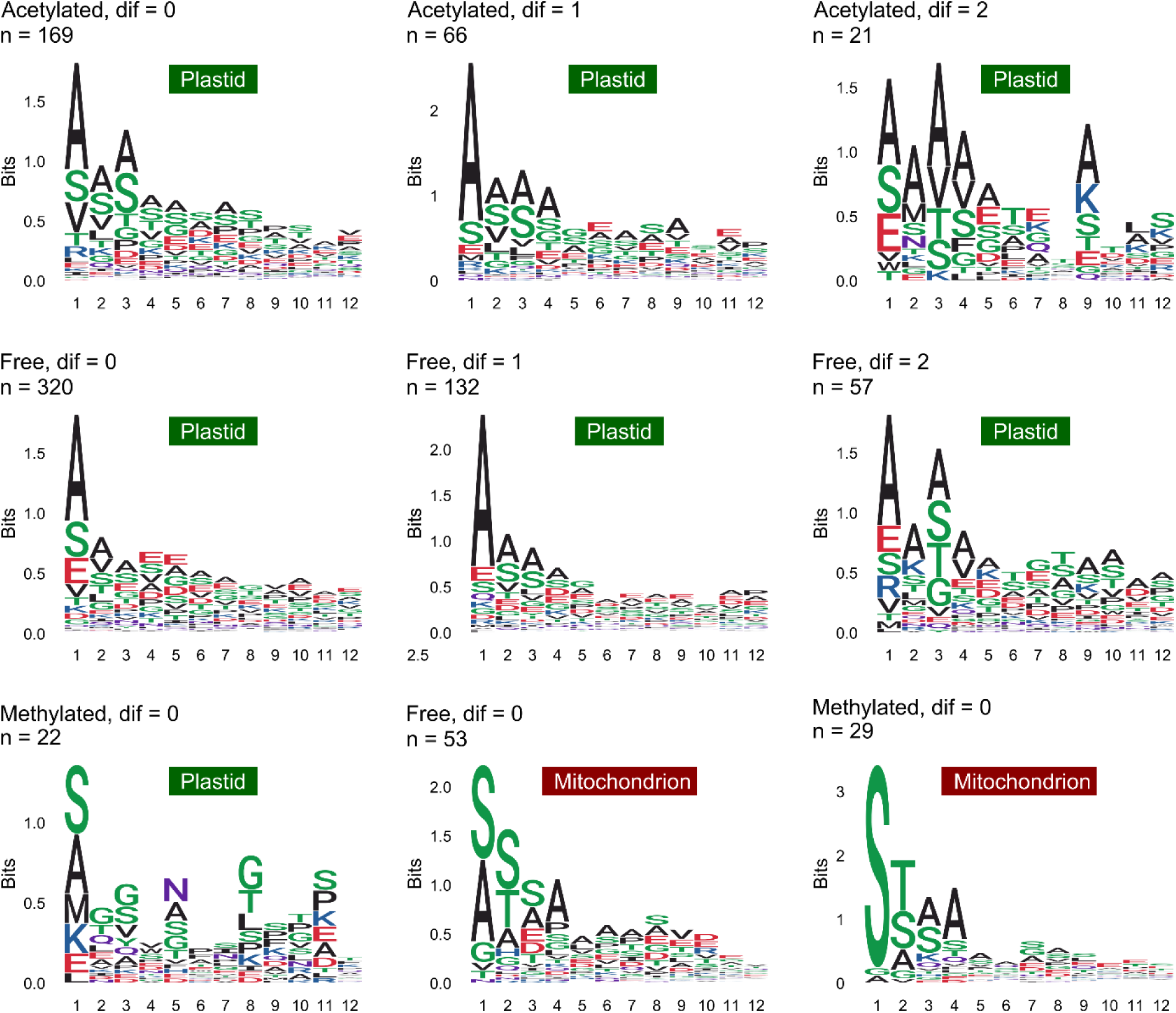
Sequence logos of identified N-termini with different modification states of plastid and mitochondrial proteins. “Dif” indicates the position difference upstream of a predicted plastid or mitochondrial transit peptide cleavage site. Transit peptide cleavage sites were predicted with *TargetP2.0*. The prediction data are available from Supplemental Table S4. “n” represents the total number of non-redundant sequences. Sequences were aligned at the identified N-terminal amino acid.

On the other hand, a cleavage of an N^α^-acetylated amino acid may also be possible, although until now no protease has clearly proven activity on acetylated N-termini of intact proteins. However, acylamino acid-releasing enzyme (AARE), a bifunctional serine protease (Tsunasawa et al. 1975; Fujino et al. 2000; Shimizu et al. 2003; Nakai et al. 2012; Hoernstein et al. 2023), has a proven activity on N^α^-acetylated oligopeptides and activity on intact proteins is repeatedly considered (Tsunasawa et al. 1975; Arfin and Bradshaw 1988; Adibekian et al. 2011). Moreover, one Physcomitrella AARE isoform and the Arabidopsis AARE are localized not only to the cytoplasm but also to plastids and mitochondria (Hoernstein et al. 2023). This renders AARE an interesting new candidate for the processing of plastid proteins, especially since this protease shows a strong substrate preference for Ac-Ala (Yamauchi et al. 2003; Hoernstein et al. 2023), the most frequent N^α^-acetylated amino acid of plastid proteins observed here (Figure 3).

We also identified several mitochondrial proteins that are N-terminally acetylated around a predicted transit peptide cleavage site (Supplemental Tables S4, S5). Although N^α^-acetylation of plastid proteins is well known, this modification has not yet been detected to a similar extent on mitochondrial proteins, and its apparent presence is not clear (Giglione and Meinnel 2021). The N-terminally acetylated amino acids were A, T, S, and in all cases were preceded by a methionine, which may also indicate alternative translation initiation. Consequently, we did not consider this to be a previously undiscovered modification of mitochondrial proteins, but rather a cotranslational modification of a shorter, e.g. cytoplasmic isoform, resulting from alternative translation initiation or alternative splicing. In Physcomitrella, both mechanisms are known to target protein isoforms to distinct subcellular localizations (Kiessling et al. 2004; Hoernstein et al. 2023). Apart from these two scenarios, dual targeting of proteins to plastids and mitochondria *via* ambiguous targeting signals is also considerable. In this case, the N-terminally acetylated protein would represent the plastid-localized variant. Hence, we investigated those N-terminally acetylated and potentially mitochondria-localized proteins with a focus on alternative translation initiation sites or splice variants that would give rise to shorter, possibly cytoplasmic, protein isoforms (Supplemental Table S5). Subcellular targeting predictions were performed with *Localizer* and the presence of ambiguous targeting signals was predicted with *ATP2*. In two cases (Pp3c13_17110V3.1, Pp3c21_2600V3.1) potential dual targeting to plastids and mitochondria was predicted by *Localizer* and *ATP2* but alternative translation initiation from the downstream methionine was also likely (Supplemental Table S5). In most other cases, we found either a potential alternative translation initiation site (Pp3c15_21480V3.1, Pp3c4_3210V3.1) or alternative splice variants (Pp3c22_8300V3.1, Pp3c7_24050V3.2) that facilitate translation of a shorter open reading frame. For one protein (Pp3c9_14150V3.1, Pp3c9_14150V3.2) the situation remains unclear. Nevertheless, the present data indicate that the observed N-terminally acetylated proteins are localized in plastids or the cytoplasm rather than in mitochondria.

Besides acetylation, we also observed N-terminal methylation on plastid proteins and on mitochondrial proteins. Strikingly, monomethylation on plastid proteins was also identified predominantly not only on N-terminal alanine and serine but also on N-terminal methionine (Figure 3). The apparent absence of a successive cleavage pattern similar to that observed for free or acetylated N-termini may indicate a stabilizing effect of monomethylation on the modified protein. This modification is found in prokaryotes and eukaryotes (Stock et al. 1987) but is poorly investigated in plants. It has been proven for the small subunit of Rubisco (RbcS) in pea, spinach, barley and corn (Grimm et al. 1997). Accordingly, we found the N-terminal methionine of RbcS to be monomethylated at its N-terminal methionine (after transit peptide removal, Pp3c12_19890V3.4, Supplemental Figure S3) in Physcomitrella, suggesting that this modification of RbcS is evolutionary conserved.

Apart from those plastid proteins, we also identify 49 mitochondrial proteins being monomethylated at their N-terminus, matching the predicted presequence cleavage site (Supplemental Table S4), including cytochrome C subunit 5B (COX5B, Pp3c19_11870V3.1, Figure S4). A strong overrepresentation of serine as N-terminally monomethylated amino acid was observed (Figure 3), whereas alanine and serine were equally frequent on non-modified N-termini of mitochondrial proteins. This specificity of methylated proteins in plastids and mitochondria resembles only partially the specificity of human NTM1A (Alpha N-terminal protein methyltransferase 1A; UNIPROT: Q9BV86) (Schaner Tooley et al. 2010; Wu et al. 2015) which methylates N-terminal alanine and serine when followed by a proline and a lysine. In our data, proline and lysine were not frequently observed as subsequent amino acids (Figure 3). Intriguingly, the observed amino acid frequency of methylated plastid and mitochondrial proteins rather resembles the situation in yeast (Chen et al. 2021). Despite RbcS, N-terminal methylation of plant proteins was reported only on cytosolic and plastid ribosomal subunits and histones (Carroll et al. 2008; Webb et al. 2010). Again, the N-terminal amino acid sequences (Webb et al. 2010) share almost no homology with the methylated N-termini observed in our data. Nevertheless, we identified the cytosolic ribosomal subunit RPL19 to be methylated at its mature N-terminus (Pp3c18_14440V3.1, Figure S5), but also other likely cytosolic proteins (Supplemental Table S3).

Finally, we also identified several proteins that were methylated at their N-terminus after cleavage of a predicted secretory signal peptide (Supplemental Table S4). Most seemed to be false positive identifications due to isotope peak errors and were not considered further.

Knowledge of N-terminal methylation of proteins, especially in plants, is scarce, and until now the methylation of RbcS was considered an exception (Grimm et al. 1997; Petkowski et al. 2013). In contrast, our data reveal that this modification affects several plastid and mitochondrial proteins with similar specificity of the methylating enzyme. In humans, two N-terminal methyltransferases are known so far, NTM1 and NTM2 (Schaner Tooley et al. 2010; Petkowski et al. 2013). To investigate whether homologues in Physcomitrella exist, we used the sequence of human NTM1 (UNIPROT: Q9BV86) as a query for a *BlastP* search (Altschul et al. 1997) against all Physcomitrella V3.3 protein models (Lang et al. 2018) using the Phytozome database (https://phytozome-next.jgi.doe.gov) (Goodstein et al. 2012) and identified a single protein (Pp3c22_8670V3.1; identity 36%, alignment length 74 amino acids) sharing the same protein family annotations as human NTM1A (InterPro: Alpha-N-methyltransferase NTM1 IPR008576; S-adenosyl-L-methionine-dependent methyltransferase IPR029063). In a reciprocal *BlastP* search using Pp3c22_8670V3.1 (hereafter referred to as PpNTM1) as a query, human NTM1 appeared as best Blast hit, confirming the orthology. A full *InterPro* search against all Physcomitrella V3.3 protein sequences (Lang et al. 2018) did not reveal any further hits with this protein family annotation. Consistent with this, PpNTM1 was also the only Blast hit when using the sequence of NTM2 (UNIPROT: Q5VVY1), a human homologue of NTM1, as a Blast query.

In Physcomitrella, PpNTM1 is expressed in all major tissues at moderate levels (Supplemental Figure S6). We also found only a single Arabidopsis homologue (AT5G44450.1) which is predicted to localize to plastids by both predictors. Interestingly, PpNTM1 is predicted via *TargetP2.0* to localize to mitochondria, whereas plastid localization is predicted by *Localizer* (Supplemental Table S6). Moreover, a potential alternative translation initiation site might be at M^56^ (Kozak Similarity Score ≥ 0.7 and < 0.8; Gleason et al. 2022a, b) which would not interfere with the predicted domain structure and enable cytosolic localization. We further investigated the conservation of residues with known catalytic function in the human isoform (Dong et al. 2015; Wu et al. 2015) in plant homologues with a focus on bryophytes, and mosses in particular, including two other species from the same family as Physcomitrella. Surprisingly, only one of seven known catalytic sites is conserved in Physcomitrella (Supplemental Figure S7A), while all but one are conserved in Arabidopsis. Notably, Arabidopsis NMT1 has a three amino acid long “EPV” motif where the human isoform has the motif “DIT” (Supplemental Figure S7A). In turn, the EPV motif appears to be conserved in all other plant species analyzed here, except the chlorophytic alga *Volvox carteri* which features an S instead of V, and Physcomitrella which deviates completely, even from the orthologues of its closest relatives (Supplemental Figure S7A). Thus, we performed phylogenetic reconstruction of the aligned protein sequences by calculating a maximum likelihood tree (Supplemental Figure S7B). The phylogeny further indicates that the accumulation of changes in Physcomitrella NTM1 are species-specific. Finally, we checked the structure predictions from AlphaFold (Jumper et al. 2021; Varadi et al. 2024). While human and Arabidopsis NTM1 have obvious structural similarities (Supplemental Figure S7C), the predicted structure of Physcomitrella NMT1 is different but of poor prediction quality (Supplemental Figure S7C). Nevertheless, the search for similar structures of PpNTM1 with *Foldseek* (van Kempen et al. 2023) again revealed sequences of NTM1 isoforms from other species such as rice (UNIPROT: Q10CT5).

Therefore, it is not yet entirely clear whether PpNTM1 is a methyltransferase responsible for the monomethylation observed here, and whether, at least in Physcomitrella, it could be targeted to both plastids and mitochondria. The present data do not provide evidence for predicted transit peptide cleavages or alternative translation initiation for this protein. Hence, further research is required to investigate the molecular function and localization of PpNTM1. Two scenarios are currently conceivable: i) The deviant Physcomitrella NMT1 is responsible for the monomethylation of N-termini observed here. A targeted gene ablation based on highly efficient homologous recombination (Hohe et al. 2004) would result in knockout mutants with no or drastically reduced monomethylated N-termini. ii) In addition to NMT1, at least one other enzyme is responsible for the observed monomethylation of N-termini in Physcomitrella and possibly also in other plants.

## Conclusion

In the present study, we used a compilation of 24 proteomic datasets obtained from different experiments to gain first insights into the N-terminome of the model plant Physcomitrella. We found that the percentage of N-terminal acetylation of cytosolic proteins appears slightly lower than the estimated percentage in Arabidopsis. Our data allow the confirmation of hundreds of predicted targeting peptide cleavage sites localizing proteins to plastids. These data can now be used to optimize computational targeting predictors, for customized protein fusions and their targeted localization in biotechnology, and provide new insights into the potential dual targeting of proteins. Furthermore, we show that N-terminal monomethylation is a previously unknown modification of mitochondrial proteins. The function and effects of this modification need to be further analyzed, but we propose PpNTM1 as a candidate for protein methylation in plastids, mitochondria and the cytosol.

## Methods

### Cell culture

For all experiments, Physcomitrella WT (new species name: *Physcomitrium patens* (Hedw.) Mitt.; Medina et al. 2019) ecotype “Gransden 2004” available from the international Moss Stock Center (IMSC, www.moss-stock-center.org, #40001) was used. Cultivation was performed using Knop medium (Reski and Abel 1985) containing 250 mg/l KH_2_PO_4_, 250 mg/l KCl, 250 mg/l MgSO_4_ x 7 H_2_O, 1,000 mg/l Ca(NO_3_)_2_ x 4 H_2_O and 12.5 mg/l FeSO_4_ x 7 H_2_O (pH 5.8). According to Egener et al. (2002) and Schween et al. (2003) 10 mL of a microelement solution 309 mg/l H_3_BO_3_, 845 mg/l MnSO_4_ x 1 H_2_O, 431 mg/l ZnSO_4_ x 7 H_2_O, 41.5 mg/l KI, 12.1 mg/l Na_2_MoO_4_ x 2 H_2_O, 1.25 mg/l CoSO_4_ x 5 H_2_O, 1.46 Co(NO_3_)_2_ x 6 H_2_O) was added per liter of medium. Gametophores were either cultivated on plates containing 12 g agar per liter liquid medium or on hydroponic ring cultures as described in Erxleben et al. (2012) and Hoernstein et al. (2016). Hydroponic gametophore cultures were started from protonema culture that was dispersed weekly using an Ultra Turrax (IKA, Staufen, Germany) at 18,000 rpm for 90 s. All cultivation was done at 25°C in a day/night cycle of 16 h light with a light intensity of 70 µmol/sm^2^ and 8 h dark.

### Treatments

Treatment with the proteasome inhibitor epoxomicin (Peptide Institute Inc., Osaka, Japan) was done using gametophores cultivated on agar plates. Gametophores were harvested and incubated in 10 mL Knop medium containing 20 µM epoxomicin for 24 h (enrichment II, Supplemental Table S1). Red-light treatment (enrichment III, Supplemental Table S1) was done using hydroponic gametophore cultures. Cultures were incubated for three days in a red-light chamber at 650 nm. Additionally, 50 µM of the proteasome inhibitor MG132 (Selleckchem, Houston, TX, USA) were applied in the culture medium at the beginning of the treatment. Dark treatments (enrichment V and VI, Supplemental Table S1) were done by wrapping the entire boxes of hydroponic gametophore cultures in aluminum foil for the indicated time and wrapped boxes were cultivated further at the same conditions as before. Proteasome inhibition of gametophores during dark treatment (enrichment VI, Supplemental Table S1) was done by submerging a hydroponic ring culture entirely in Knop medium containing 100 µM MG132. The box was wrapped in aluminum foil and incubated for 24 h.

### Enrichment of nuclei from gametophores

Eighteen g fresh weight (FW) gametophores were harvested from hydroponic culture and chopped in buffer I containing 1 M 2-Methyl-2,4-pentandiol, 10 mM HEPES pH 7.5, 10 mM KCl 10 mM DTT, 0.1% PVP40, 0.1% PPI (P9599, Sigma-Aldrich, St. Louis, MO, USA) according to Nelson et al. (1994) using a custom 4 razorblade chopping device. The homogenate was successively filtered through a 40 µm and a 20 µm sieve and the flow-through was centrifuged for 30 min at 300 x g at 2°C. The supernatant was discarded, and the pellets were carefully dissolved in buffer II containing 110 mM KCl, 15 mM HEPES, pH 7.5, 5mM DTT and 0.1% PPI. The enriched nuclei were further purified using three-step Percoll-gradients (100%/60%/30%, 17-0891-01, GE Healthcare, Solingen, Germany) modified after Marienfeld et al. (1989). The Percoll-gradients were centrifuged at 200 x g at 2°C for 30 min. The interface between 100% and 60% was recovered as well as the pellet at the top of the gradient attached to the tube wall. Both fractions were strongly enriched in nuclei and thus pooled for further experiments. The samples were combined and washed with buffer II and centrifuged again for 10 min at 300 x g at 2°C. The pellet containing enriched nuclei was stored at −20°C until further use.

### Sequential protein extraction from nuclei

Pellets containing enriched nuclei were dissolved in 400 µL 50 mM Tris-HCl, pH7.6, 1% PPI (P9599, Sigma-Aldrich) and sonicated (Sonopuls HD2070, Bandelin, Berlin, Germany) three times for 20 s with an amplitude between 60 and 90%. Fifty µL DNAse buffer and 50 µL DNase I (EN0521, Thermo Scientific, Waltham, USA) were added and the samples were incubated for 1 h at 37°C. After centrifugation at 20,000 x g for 20 min at 4°C the supernatant (Tris extract) was recovered and directly precipitated with 5 vol ice-cold acetone containing 0.2% DTT overnight at −20°C. The remaining pellet was dissolved in 400 µL 50 mM Tris-HCl, pH 7.6, 2% Triton X-100, 1% PPI, and again sonicated three times as before. Again, the sample was centrifuged, and the supernatant (Triton extract) was also acetone-precipitated overnight. The remaining pellet was dissolved in 50 mM Tris-HCl, pH 7.6, 4% SDS, 1% PPI, 50 mM DTT and incubated at 95°C for 10 min. The sample was centrifuged, and the supernatant was acetone-precipitated overnight. All acetone precipitations were centrifuged at 20,000 x g at 0°C for 15 min. The supernatant was discarded, and the remaining protein pellet was washed for 1 h with 1 vol ice-cold acetone without DTT. The centrifugation step was repeated, and the supernatant was discarded afterwards. The remaining protein pellets were air dried and stored at −20°C for further experiments.

### Sequential protein extraction from gametophores

One to two g FW of gametophores were ground in liquid nitrogen for 10-15 min. The fine powder was dissolved in Tris buffer containing 40 mM Tris-HCl, pH 7.6, 0.5% PVPP and 1% PPI. The homogenate was sonicated for 15 min and afterwards centrifuged at 20,000 x g at 4°C for 30 min. The supernatant (Tris-extract) was recovered. The remaining pellet containing cell debris was dissolved in 40 mM Tris-HCl pH 7.6, 2% Triton X-100, 1% PPI and again sonicated for 15 min. Again, centrifugation was performed at 20,000 x g at 4°C for 30 min and the supernatant (Triton-extract) was recovered. Protein concentrations of the extracts were directly determined via the Bradford assay (Bradford 1976) and aliquots corresponding to 100 µg protein were precipitated with acetone containing DTT as described before.

### Enrichment of N-terminal peptides

The dimethylation reaction was carried out according to Kleifeld et al. (2010) with some modifications. Protein pellets were dissolved in 100 mM HEPES-NaOH pH 7.5, 0.2% SDS. Reduction of cysteine residues was carried out using *Reducing Agent* (NP0009, Life Technologies™, Carlsbad, USA) 1:10 at 95°C for 10 min or *Bond-Breaker®* (77720, Thermo Scientific) 1:100 at 28°C for 30 min. Alkylation was performed at a final concentration of 100 mM iodoacetamide for 20 min at RT. The dimethylation reaction was carried out by adding 2 µL of a 4% formaldehyde solution (Formaldehyde ^13^C, d_2_ solution, 596388, Sigma-Aldrich or Formaldehyde-D2, DLM-805-PK, Cambridge Isotope Laboratories Inc.) and 2 µL of a 500 mM NaCNBH_3_ solution per 100 µL sample at 37°C for 4 h. The same volumes of formaldehyde and NaCNBH_3_ were added again to the sample and the reaction was carried out overnight at 37°C. The dimethylation reaction was stopped by adding 2 µL of a 4% NH_4_OH solution per 100 µL sample for 1 h at 37°C. Afterwards the samples were precipitated as described before using acetone without DTT for at least 3 h at −20°C. The final enrichment was modified according to McDonald and Beynon (2006).

### SDS-based enrichment

The dried protein pellets were dissolved in binding buffer according to McDonald and Beynon (2006) containing 20 mM NaH_2_PO_4_, 150 mM NaCl pH 7.5 with 0.2% SDS and in solution digest using either trypsin (V5280, Promega, Madison, USA), GluC (90054, Thermo Scientific) or chymotrypsin (V1062, Promega) was performed at an enzyme-to-substrate ratio of 1:25 for 4 h at 37°C (trypsin, GluC) or 25°C (chymotrypsin). Then the ratio was increased to 1:20 and the reaction was carried out overnight. Enrichment of N-terminal labeled peptides was carried out using 200 µL NHS-sepharose slurry (17-0906-01, GE Healthcare, Solingen, Germany) per 100 µg protein. The slurry was centrifuged for 30 sec at 200 x g. The supernatant was discarded and 400 µL ice-cold 1 mM HCl was added. The slurry was centrifuged again, and the supernatant was discarded. Afterwards the sepharose was washed with 1 mL binding buffer without SDS. The samples were applied to the prepared sepharose and incubated for 4 h at RT. The sepharose was again centrifuged, and the supernatant was transferred to a new tube containing freshly prepared sepharose. The used sepharose was washed with 20 µL binding buffer and the supernatant was also added to the freshly prepared sepharose. The enrichment reaction was carried out overnight at 4-8°C. The enriched peptides were desalted using 200 µl C18 *StageTips* (SP301, Thermo Scientific) that were supplemented with an additional layer of *Empore^TM^ SPE Disk* C18 material (66883-U, Sigma-Aldrich). The tips were washed prior to use with 100 µl 0.1% TFA and subsequently with 100 µl 80% ACN, 0.1% TFA. The tips were again equilibrated with 100 µl 0.1% TFA and the samples were loaded afterwards. The remaining sepharose was washed with 50 µl binding buffer and the supernatant was also transferred to the tip. The tips were washed with 100 µl binding buffer and the retained peptides were eluted with 300 µl 80% ACN, 0.1% TFA. The eluate was vacuum dried and the samples were stored at −20°C until further analysis.

### *Rapi*Gest-based enrichment

The dried protein pellets were dissolved in 50 mM HEPES-KOH, 0.1% *Rapi*Gest surfactant (RPG, 18600186, Waters, Milford Massachusetts, USA). Proteolytic digest was performed as described before. After digestion, the RPG surfactant was cleaved by acidifying the sample to pH 2 using TFA as recommended by the manufacturer. The cleavage was performed at 37°C for 45 min. Insoluble RPG remnants were removed by centrifugation at 13,000 rpm for 10 min at RT. The peptide-containing supernatant was subjected to solid phase extraction using SampliQ C18 cartridges (1 ml, 100 mg, 5982-1111, Agilent, Santa Clara, USA). Prior to the extraction the cartridge was washed successively with 1 ml 0.1% TFA,0.1% TFA in 80% ACN, 0.1% TFA. Then the peptide solution was applied and washed with1 ml 0.1% TFA. The peptides were eluted in 600 µl 0.1% TFA in 80% ACN and vacuum dried. The dried peptides were stored at −20°C until further use. For enrichment of the N-terminal peptides, the dried and purified peptides were dissolved in binding buffer as described before without SDS and the enrichment using NHS-sepharose was performed accordingly. Finally, the enriched N-terminal peptides were desalted again using SampliQ C18 cartridges as described before. The dried peptides were stored at −20°C until mass spectrometric analysis.

### Mass spectrometry

NanoLC-MS/MS analyses were performed on an LTQ-Orbitrap Velos Pro (ThermoScientific) equipped with an EASY-Spray Ion Source and coupled to an EASY-nLC 1000 (Thermo Scientific). Peptides were loaded on a trapping column (2 cm x 75 µm ID. PepMap C_18_ 3 µm particles, 100 Å pore size, Dionex, Thermo Scientific) and separated either on a 25 cm EASY-Spray column (25 cm x 75 µm ID, PepMap C_18_ 2 µm particles,100 Å pore size) with a 30 min linear gradient from 3% to 30% ACN (V5280, Promochem) and 0.1% FA (56302, Thermo Scientific), or on a 50 cm EASY-Spray column (50 cm x 75 µm ID, PepMap C_18_ 2 µm particles, 100 Å pore size, Dionex, Thermo Scientific) with a 360 min linear gradient from 3% to 30% ACN and 0.1% FA in the case of in solution digested proteins such as the enriched N-terminal peptides. MS scans were acquired in the Orbitrap analyzer with a resolution of 30,000 at m/z 400, MS/MS scans were acquired in the Orbitrap analyzer with a resolution of 7500 at m/z 400 using HCD fragmentation with 30% normalized collision energy. A TOP5 or TOP10 data-dependent MS/MS method was used. Dynamic exclusion was applied with a repeat count of 1 and an exclusion duration of 30 sec or 2 min in the case of long gradients. Singly charged precursors were excluded from selection. Minimum signal threshold for precursor selection was set to 50,000. Predictive AGC was used with a target value of 10^6^ for MS scans and 5*10^4^ for MS/MS scans. Lock mass option was applied for internal calibration using background ions from protonated decamethylcyclopentasiloxane (m/z 371.10124). Electron-transfer dissociation (ETD) fragmentation was performed with 35% normalized collision energy. A TOP5 data dependent MS/MS method was used. Dynamic exclusion was applied with a repeat count of 1 and an exclusion duration of 30 seconds. Singly charged precursors were excluded from selection. Minimum signal threshold for precursor selection was set to 75,000. Predictive AGC was used with AGC target a value of 10^6^ for MS scans and 5*10^4^ for MS/MS scans. ETD activation time was set to 250 ms for doubly charged precursors, 166 ms for triply charged precursors and 125 ms for quadruply charged precursors, and the AGC target was set to 200,000 for fluoranthene. Lock mass option was applied for internal calibration in all runs using background ions from iron(III) citrate (m/z 263.956311).

### Raw data processing and database search

Raw data were processed with *Mascot Distiller* (V2.8.3.0, https://www.matrixscience.com/) and database searches were performed using *Mascot Server* (V2.7.0, https://www.matrixscience.com) against a database containing all V3 Physcomitrella protein models (Lang et al. 2018) as well as their reversed sequences as decoys. In parallel, a search was performed against a database containing the sequences of known contaminants, such as keratin (269 entries, available on request). For all samples semi-specific protease specificities were chosen and in the case of tryptic digests the specificity was set to semi-ArgC. Variable modifications were Gln >pyro Glu (N term Q) −17.026549 Da, oxidation (M) +15.994915 Da, acetyl (N-term) +42.010565 Da, ^13^C,D_2_ dimethyl (N-term) +34.063117 Da, hybrid-methylation (N-term) +31.047208 (^13^CD_2_CH_2_) Da and deamidation of asparagine (N) +0.984016 Da. Fixed modifications were carbamidomethyl (C) +57.021464 Da and ^13^C,d_2_ dimethyl (K) +34.063117 Da. In the case of samples dimethylated with D_2_ formaldehyde, the dimethylation (K, N-term) mass shift was +32.056407 Da and hybrid-methylation (N-term) was set to +30.043854 Da (C_2_D_2_H_2_). A precursor mass tolerance of ±8 ppm and a fragment mass tolerance of ±0.02 Da was specified. Search results were loaded in *Scaffold5^TM^* software (V5.0.1, https://www.proteomesoftware.com/) using the high mass accuracy scoring and independent sample grouping method.

### Computational analysis

The presence of cleavable N-terminal targeting signals (plastid, mitochondrion, secretome) was performed with *TargetP2.0* (Armenteros et al. 2019) and in selected cases with *Localizer* (Sperschneider et al. 2017). Ambiguous targeting to plastids and mitochondria in Physcomitrella was predicted with *ATP2* (Fuss et al. 2013). Potential alternative translation initiation was predicted with TIS (https://www.tispredictor.com/) (Gleason et al. 2022a, b). All plots and tables were created using custom PERL scripts and *R* (R Core Team 2024).

### Multiple sequence alignment and phylogenetic reconstruction

The amino acid sequence of human NTM1 (UNIPROT: Q9BV86) was used as the query in BLASTP-like searches with DIAMOND (Buchfink et al. 2021) in “ultra-sensitive” mode against the proteomes of *Anthoceros angustus* (Zhang et al. 2020), *Amborella trichopoda* (Amborella Genome Project et al. 2013), *Arabidopsis thaliana* (Cheng et al. 2017), *Calohypnum plumiforme* (Mao et al. 2020), *Ceratodon purpureus* (Carey et al. 2021), *Funaria hygrometrica* (Kirbis et al. 2020), *Marchantia polymorpha* (Bowman et al. 2017), *Oryza sativa* (Ouyang et al. 2007), Physcomitrella (Lang et al. 2018), *Physcomitrellopsis africana* (Vuruputoor et al. 2024), *Selaginella moellendorffii* (Banks et al. 2011), *Sphagnum fallax* (Healey et al. 2023), *Sphagnum magellanicum* (Healey et al. 2023), *Takakia lepidozioides* (Hu et al. 2023) and *Volvox carteri* (Prochnik et al. 2010). Best hits were validated by reciprocal BLAST and aligned with MAFFT (Katoh and Standley 2013) in “localpair” mode with a maximum of 1000 iterations. Phylogenetic reconstruction was performed using RAxML-NG (Kozlov et al. 2019) using the ‘JTT-DCMUT+G4’ model and 1000 bootstrap replicates, rooted at the split between human and plants and visualized using R (R Core Team 2024) and the ggtree package (Yu et al. 2017).

## Data availability

The mass spectrometry proteomics data have been deposited to the *ProteomeXchange* Consortium via the *PRIDE* partner repository (Perez-Riverol et al. 2022; Deutsch et al. 2023) with the dataset identifier PXD052824 and 10.6019/PXD052824.

## Author contributions

R.R., G.L.I. and A.S. designed research and acquired funding. D.L. supervised research and helped analyzing the data. S.N.W.H. and K.F. performed experiments. N.v.G. analyzed data. S.N.W.H. analyzed the data and wrote the manuscript together with R.R. All authors approved the final version of the manuscript.

## Acknowledgements

Funding by Deutsche Forschungsgemeinschaft (DFG IG9/8-1 to G.L.I., A.S. and R.R.) and under Germany’s Excellence Strategy (CIBSS – EXC-2189 – Project ID 390939984) and by Wissenschaftliche Gesellschaft Freiburg is gratefully acknowledged. We thank Christine Glockner and Stephanie Lamer for technical assistance and Anne Katrin Prowse for proof reading.

## Supplementary information

**Figure S1.**
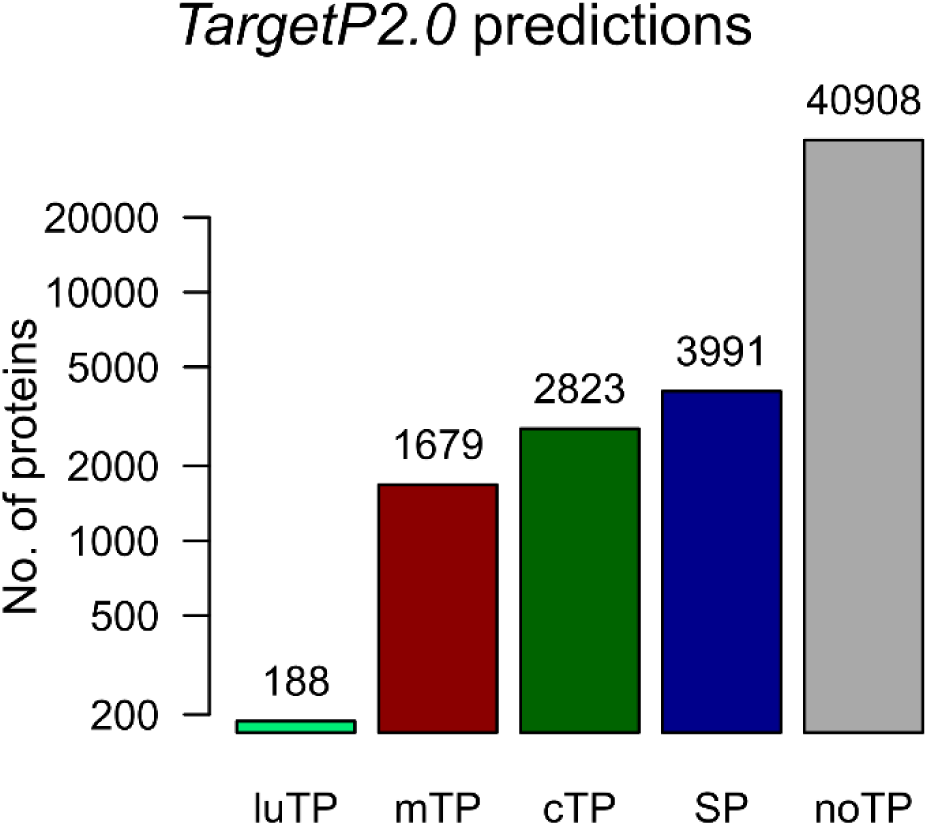
Overview on the numbers of cleavable targeting sequences in Physcomitrella predicted by *TargetP2.0*. Predictions were done on a non-redundant protein isoform list (49,589 entries) of all Physcomitrella V3.3 protein models (Lang et al. 2018). luTP: thylakoid luminal targeting peptide; mTP: mitochondrial targeting peptide; cTP: plastid targeting peptide; SP: (secretory) signal peptide; noTP: no targeting peptide predicted.

**Figure S2.**
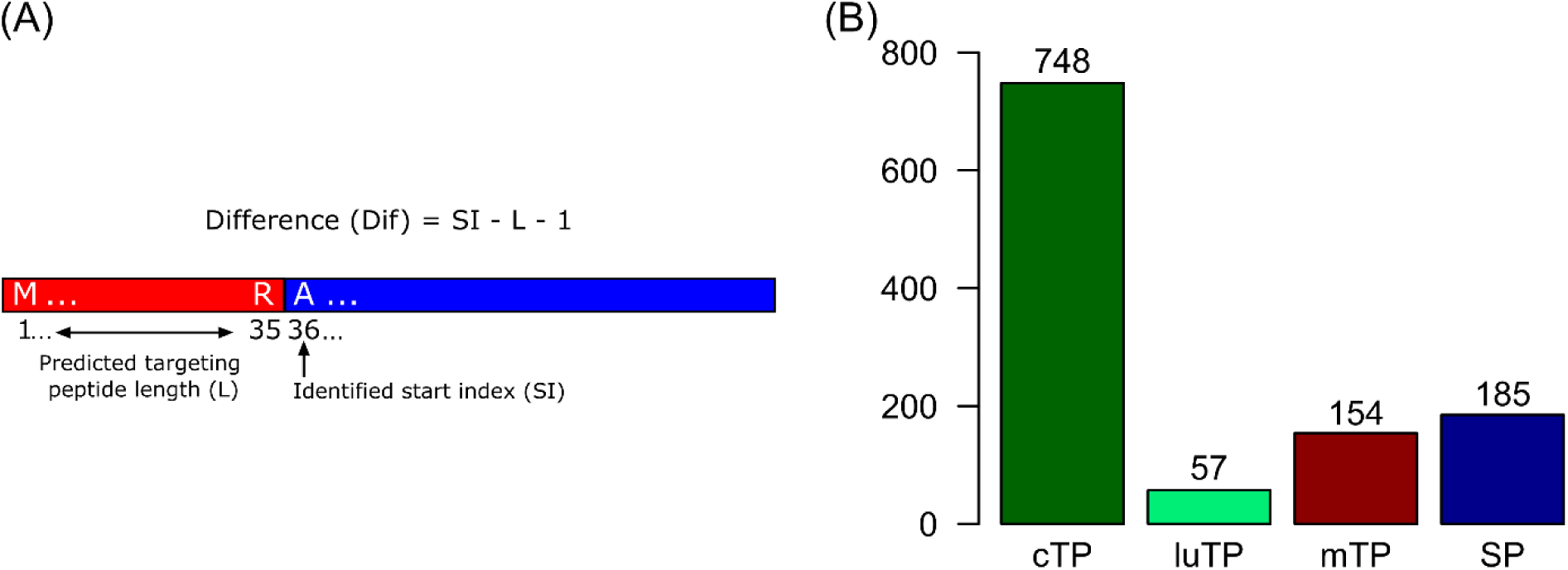
Overview on targeting peptide cleavage site identification and summary of identified targeting peptide cleavage sites. (**A**) Scheme depicting the calculation of the difference between predicted targeting peptide cleavage site and experimentally observed N-terminus. A difference of 0 indicates full agreement between prediction and observation. (**B**) Summary of identified plastid (cTP), luminal (luTP), mitochondrial (mTP) and secretory (SP) targeting peptide cleavage sites. A difference of ±5 from a predicted cleavage site was accepted. Data are available from Supplemental Table S4.

**Figure S3.**
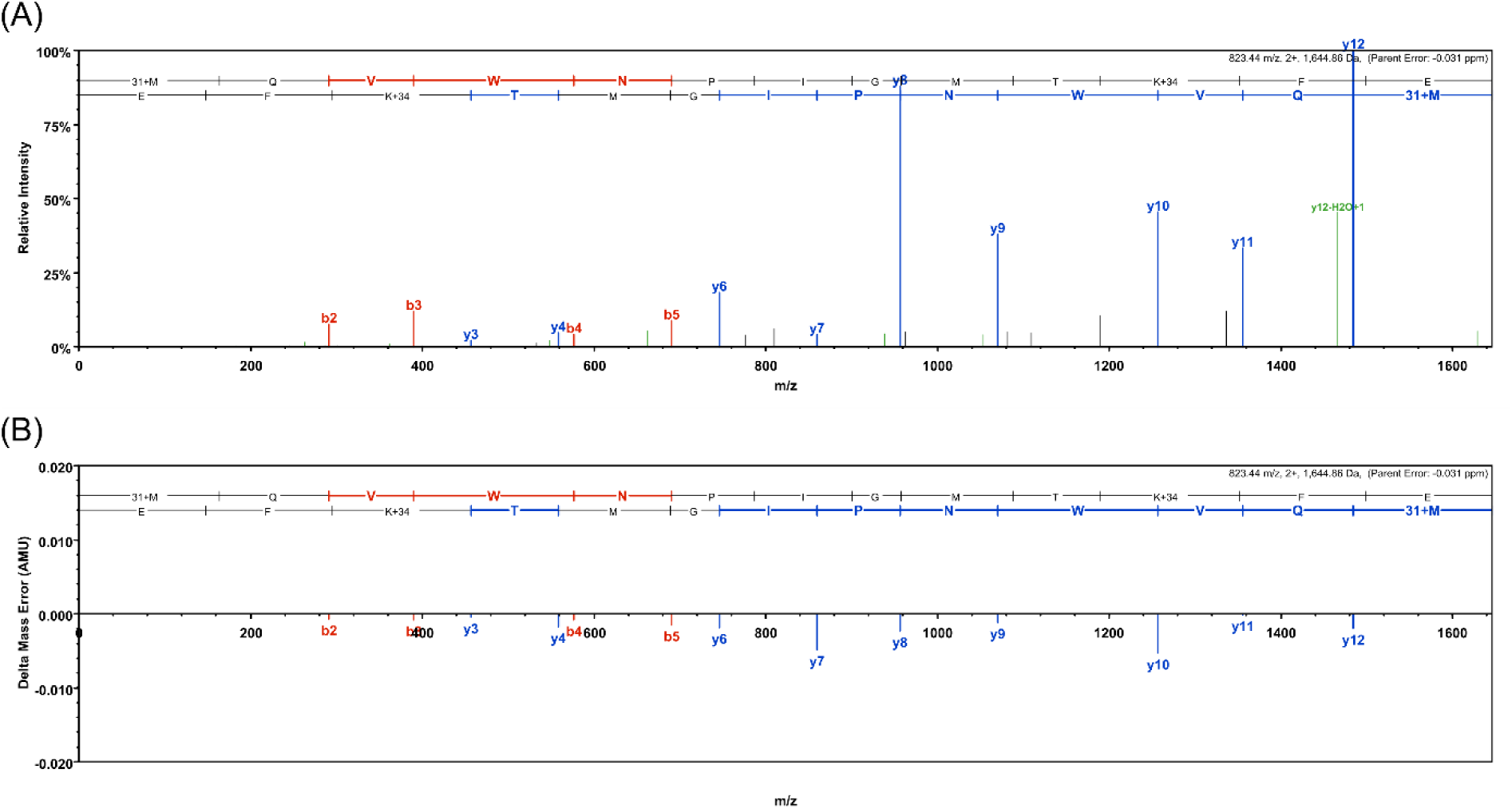
Higher-energy collisional dissociation (HCD) fragment mass spectrum and fragment mass error distribution of the identified N-terminal peptide of RbcS (Pp3c12_19890V3.4). (**A**) HCD fragment mass spectrum of the peptide MQVWNPIGMTKFE. A mass shift of +31 indicates the hybrid modification of a post-translationally incorporated methyl group and the further addition methyl group from the preformed reductive methylation during sample preparation (^13^CD_2_CH_2_, +31.047208). (**B**) Fragment mass error distribution of the b- and y-ion series.

**Figure S4.**
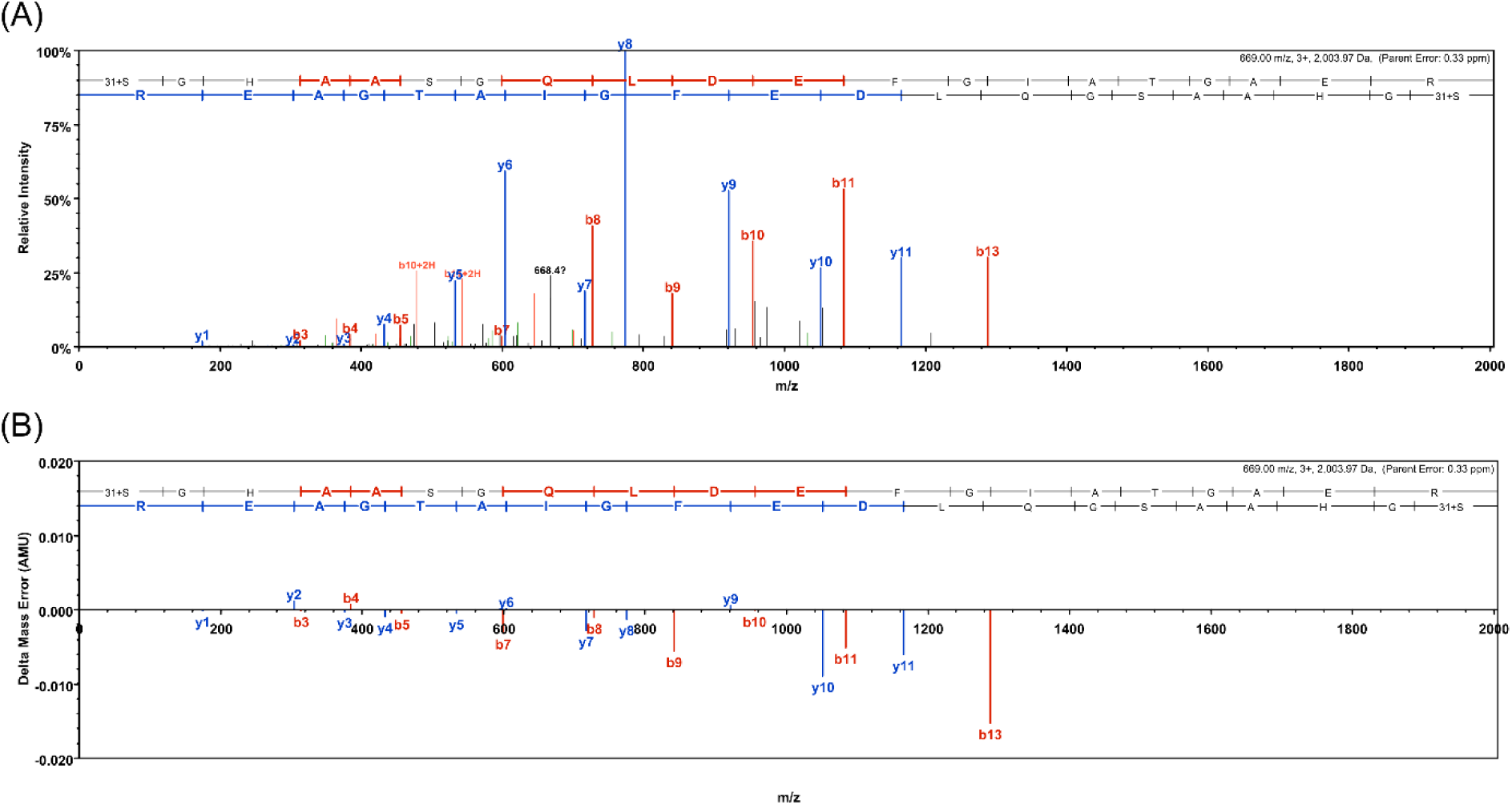
Higher-energy collisional dissociation (HCD) fragment mass spectrum and fragment mass error distribution of the identified N-terminal peptide of COX5B (Pp3c19_11870V3.1). (**A**) HCD fragment mass spectrum of the peptide SGHAASGQLDEFGIATGAER. A mass shift of +31 indicates the hybrid modification of a post-translationally incorporated methyl group and the further addition methyl group from the preformed reductive methylation during sample preparation (^13^CD_2_CH_2_, +31.047208). (**B**) Fragment mass error distribution of the b- and y-ion series.

**Figure S5.**
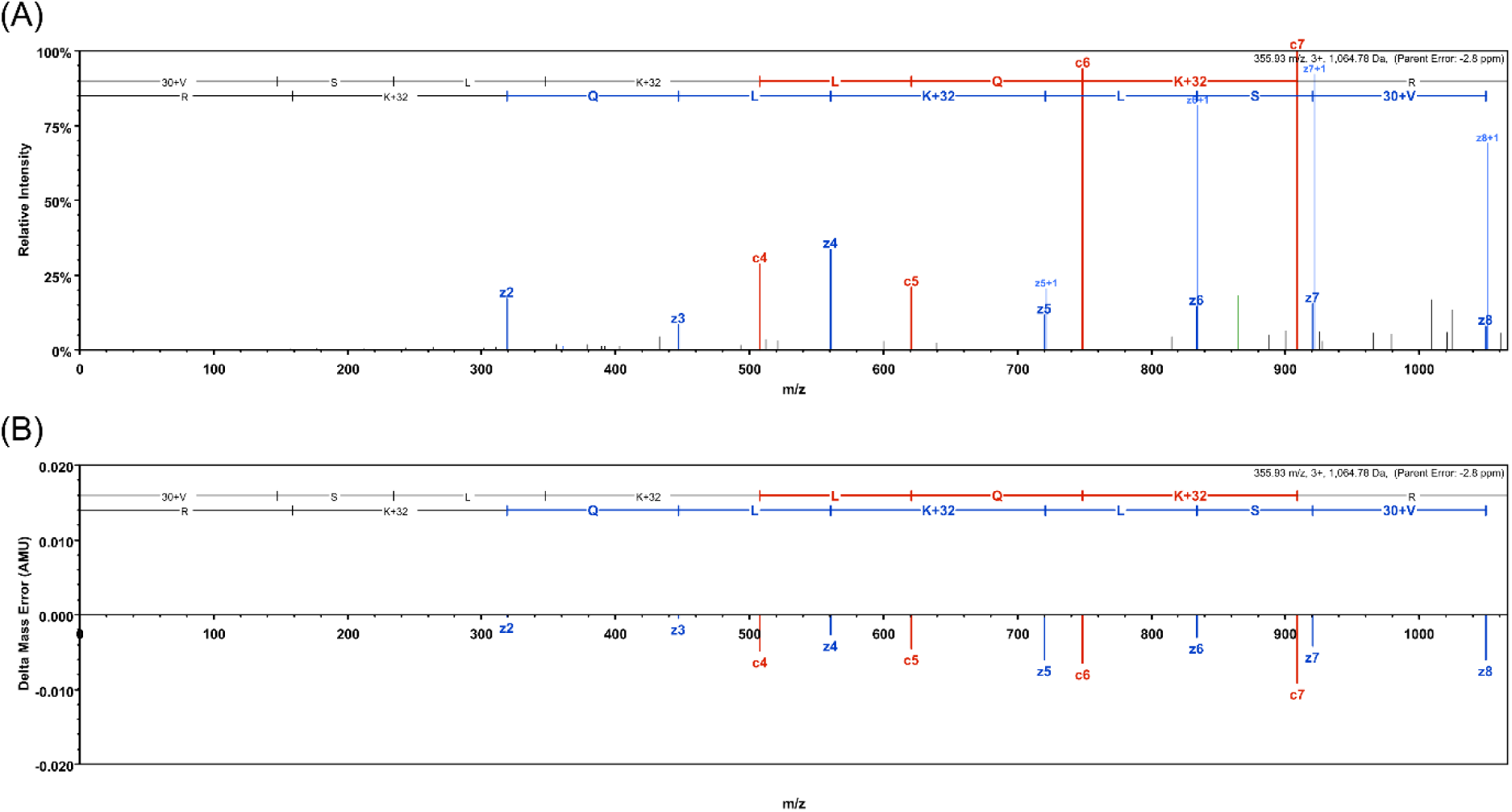
Electron-transfer dissociation (ETD) fragment mass spectrum and fragment mass error distribution of the identified N-terminal peptide of RPL19 (Pp3c18_14440V3.1). (**A**) ETD fragment mass spectrum of the peptide GKQISEIKDFLLTAR. A mass shift of +30 indicates the hybrid modification of a post-translationally incorporated methyl group and the further addition methyl group from the preformed reductive methylation during sample preparation (C_2_D_2_H_2,_ +30.043854 Da). (**B**) Fragment mass error distribution of the b- and y-ion series.

**Figure S6.**
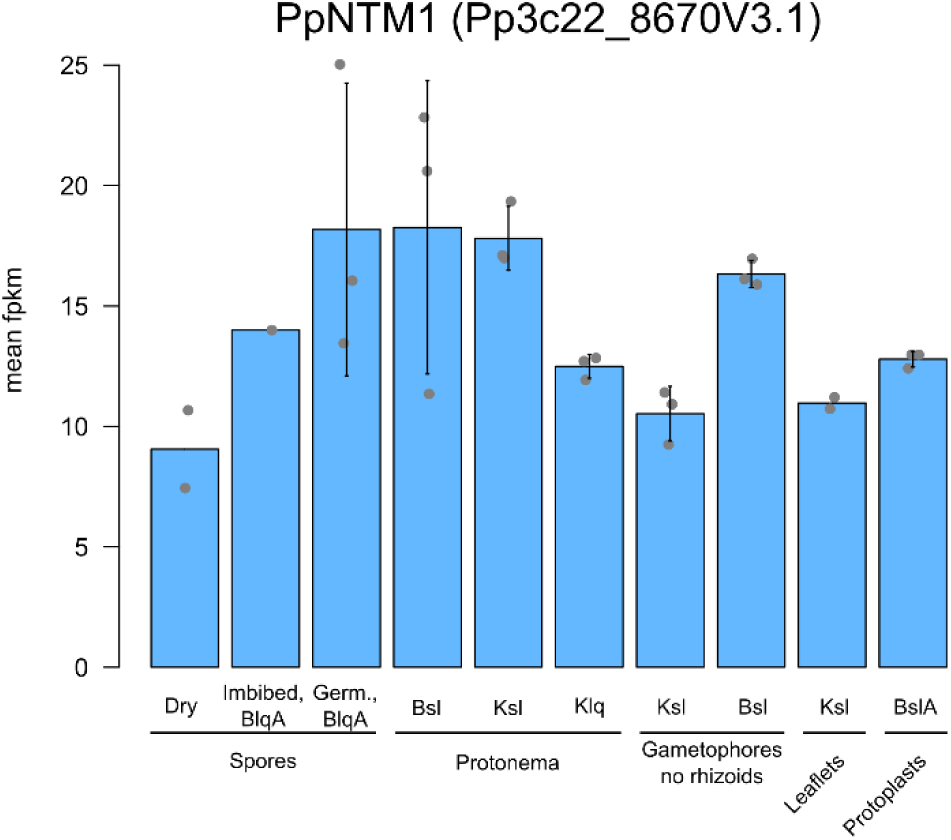
Expression levels of PpNTM1 in different tissues and cell culture types of Physcomitrella. Expression levels are represented as mean fpkm (fragments per kilobase million) values with standard deviation. Single datapoints represent biological replicates. All replicate data were downloaded from *PEATmoss* (https://peatmoss.plantcode.cup.uni-freiburg.de) and sampling and methods are described in Perroud et al. (2018) and Fernandez-Pozo et al. (2020). Abbreviations: B = BCD medium (Ashton et al. 1979); K = Knop medium (Reski and Abel 1985); A =ammonium tartrate; lq = liquid; sl = solid; Germ. = germinating.

**Figure S7.**
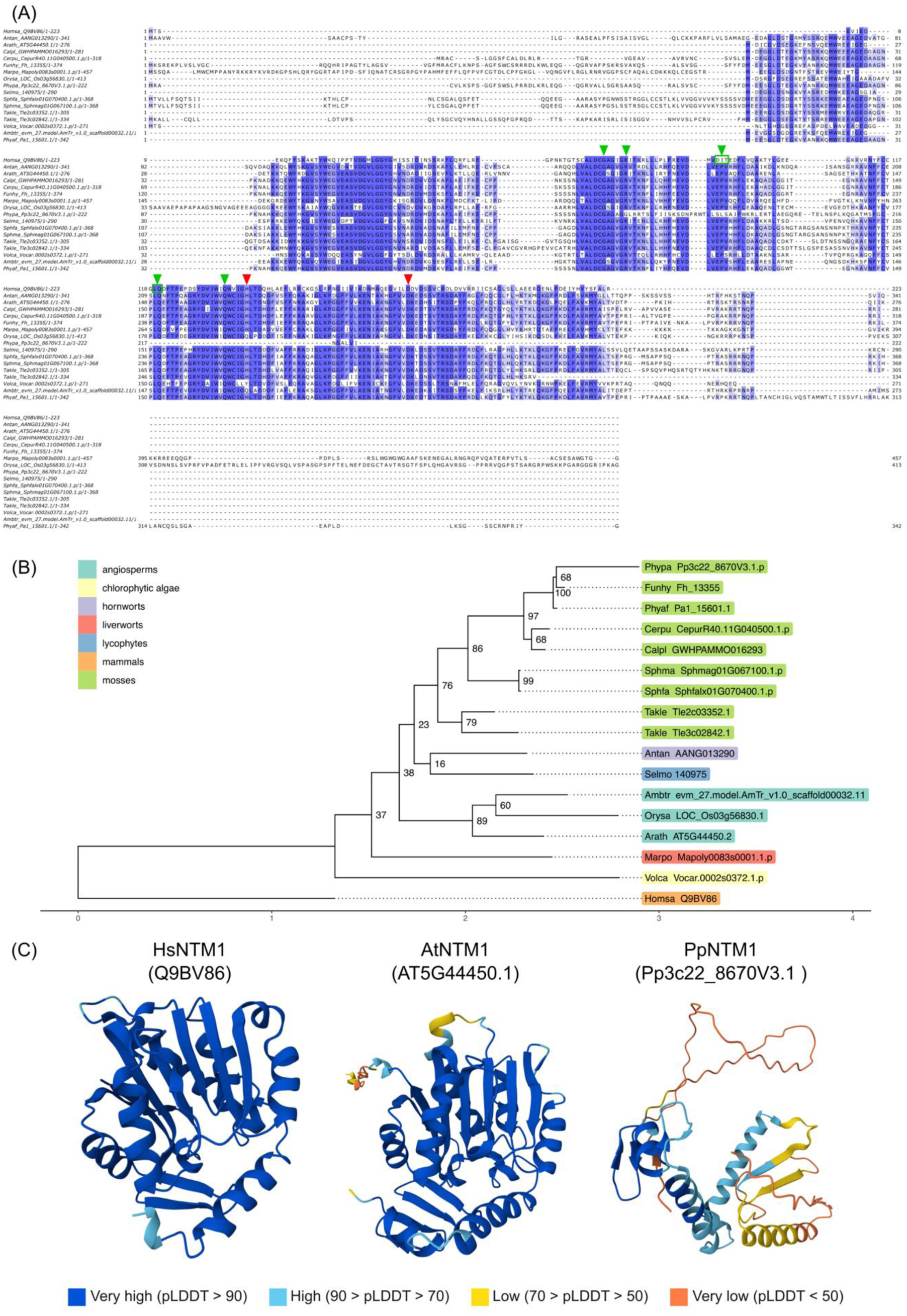
Multiple Sequence alignment of selected NTM1 sequences, phylogenetic reconstruction and AlphaFold structures. (**A**) Aligned homologous NTM1 amino acid sequences of human (Homsa), *Anthoceros angustus* (Antan), *Amborella trichopoda* (Ambtr), *Arabidopsis thaliana* (Arath), *Calohypnum plumiforme* (Calpl), *Ceratodon purpureus* (Cerpu), *Funaria hygrometrica* (Funhy), *Marchantia polymorpha* (Marpo), *Oryza sativa* (Orysa), *Physcomitrellopsis africana* (Phyaf), *Selaginella moellendorffii* (Selmo), *Sphagnum fallax* (Sphfa), *Sphagnum magellanicum* (Sphma), *Takakia lepidozioides* (Takle), *Volvox carteri* (Volca), and Physcomitrella (Phypa). Residues involved in binding of S-adenosyl-methionine (SAM) are indicated of HsNTM1 with red arrows (Dong et al. 2015; Wu et al. 2015), residues involved in target deprotonation are marked with green arrows (Dong et al. 2015; Wu et al. 2015). (B) Maximum likelihood tree of aligned NTM1 proteins with bootstrap support values at internal nodes and color-coded groups of taxa. (C) Structures of human Arabidopsis, and Physcomitrella NTM1 predicted with AlphaFold (Jumper et al. 2021; Varadi et al. 2024). Structures were accessed via UNIPROT. The prediction accuracy of the models is indicated by color representing the per-residue model confidence score (pLDDT: predicted local distance difference test; Varadi et al. 2024).

**Table S1** Overview on the performed experiments, employed culture type and sample processing

**Table S2** Protein identification summary on all experiments exported from *Scaffold5* software

**Table S3** Overview on identified N-terminal peptides

**Table S4** Comparison of identified N-terminal positions with predicted transit peptide cleavage sites

**Table S5** Overview on proteins possibly dually targeted to mitochondria and the cytosol or to plastids

**Table S6** Overview on subcellular predictions for PpNTM1

## References

Adibekian A, Martin BR, Wang C, Hsu KL, Bachovchin DA, Niessen S, Cravatt BF (2011) Click-generated triazole ureas as ultrapotent *in vivo*–active serine hydrolase inhibitors. Nat Chem Biol 7:469–478. 10.1038/nchembio.579

Altschul SF, Madden TL, Schäffer AA, Zhang J, Zhang Z, Miller W, Lipman DJ (1997) Gapped BLAST and PSI-BLAST: a new generation of protein database search programs. Nucleic Acids Res 25:3389–3402. 10.1093/nar/25.17.3389

Amborella Genome Project, Albert VA, Barbazuk WB, DePamphilis CW, Der JP, Leebens-Mack J, Ma H, Palmer JD, Rounsley S, Sankoff D et al. (2013) The *Amborella* genome and the evolution of flowering plants. Science 342:1241089. 10.1126/science.1241089

Arfin SM, Bradshaw RA (1988) Cotranslational processing and protein turnover in eukaryotic cells. Biochem 27:7979–7984. 10.1021/bi00421a001

Armenteros JJA, Salvatore M, Emanuelsson O, Winther O, von Heijne G, Elofsson A, Nielsen H (2019) Detecting sequence signals in targeting peptides using deep learning. Life Sci Alliance 2:e201900429. 10.26508/lsa.201900429

Banks JA, Nishiyama T, Hasebe M, Bowman JL, Gribskov M, dePamphilis C, Albert VA, Aono N, Aoyama T, Ambrose BA, et al. (2011) The Selaginella genome identifies genetic changes associated with the evolution of vascular plants. Science 332:960–963. 10.1126/science.1203810

Bi G, Zhao S, Yao J, Wang H, Zhao M, Sun Y, Hou X, Haas FB, Varshney D, Prigge M et al. (2024) Near telomere-to-telomere genome of the model plant *Physcomitrium patens*. Nat Plants 10:327–343. 10.1038/s41477-023-01614-7

Bienvenut WV, Sumpton D, Martinez A, Lilla S, Espagne C, Meinnel T, Giglione C (2012) Comparative large scale characterization of plant versus mammal proteins reveals similar and idiosyncratic N-α-acetylation features. Mol Cell Proteomics 11.6. 10.1074/mcp.M111.015131

Bowman JL, Kohchi T, Yamato KT, Jenkins J, Shu S, Ishizaki K, Yamaoka S, Nishihama R, Nakamura R, Berger F et al. (2017) Insights into land plant evolution garnered from the *Marchantia polymorpha* genome. Cell 171:287–304.e15. 10.1016/j.cell.2017.09.030

Bradford MM (1976) A rapid and sensitive method for the quantitation of microgram quantities of protein utilizing the principle of protein-dye binding. Anal Biochem 72:248–254. 10.1016/0003-2697(76)90527-3

Buchfink B, Reuter K, Drost HG (2021) Sensitive protein alignments at tree-of-life scale using DIAMOND. Nat Methods 18:366–368. 10.1038/s41592-021-01101-x

Carey SB, Jenkins J, Lovell JT, Maumus F, Sreedasyam A, Payton AC, Shu S, Tiley GP, Fernandez-Pozo N, Healey A et al. (2021) Gene-rich UV sex chromosomes harbor conserved regulators of sexual development. Sci Adv 7:eabh2488. 10.1126/sciadv.abh2488

Carroll AJ, Heazlewood JL, Ito J, Millar AH (2008) Analysis of the *Arabidopsis* cytosolic ribosome proteome provides detailed insights into its components and their post-translational modification. Mol Cell Proteomics 7:347–369. 10.1074/mcp.M700052-MCP200

Chen P, Paschoal Sobreira TJ, Hall MC, Hazbun TR (2021) Discovering the N-terminal methylome by repurposing of proteomic datasets. J Proteome Res 20:4231–4247. 10.1021/acs.jproteome.1c00009

Cheng C, Krishnakumar V, Chan AP, Thibaud-Nissen F, Schobel S, Town CD (2017) Araport11: a complete reannotation of the *Arabidopsis thaliana* reference genome. Plant J 89:789–804. 10.1111/tpj.13415

Decker EL, Reski R (2020) Mosses in biotechnology. Curr Opin Biotech 61:21–27. 10.1016/j.copbio.2019.09.021

Decker EL, Alder A, Hunn S, Ferguson J, Lehtonen MT, Scheler B, Kerres KL, Wiedemann G, Safavi-Rizi V, Nordzieke S et al. (2017) Strigolactone biosynthesis is evolutionarily conserved, regulated by phosphate starvation and contributes to resistance against phytopathogenic fungi in a moss, *Physcomitrella patens*. New Phytol 216:455–468. 10.1111/nph.14506

Demir F, Perrar A, Mantz M, Huesgen PF (2022) Sensitive plant N-terminome profiling with HUNTER. Methods in Molecular Biology 2247:139–158. 10.1007/978-1-0716-2079-3_12

Deutsch EW, Bandeira N, Perez-Riverol Y, Sharma V, Carver JJ, Mendoza L, Kundu DJ, Wang S, Bandla C, Kamatchinathan S et al. (2023) The ProteomeXchange consortium at 10 years: 2023 update. Nucleic Acids Res 51:D1539–D1548. 10.1093/nar/gkac1040

Dong C, Mao Y, Tempel W, Qin S, Li L, Loppnau P, Huang R, Min J (2015) Structural basis for substrate recognition by the human N-terminal methyltransferase 1. Genes Dev 29:2343–2348. http://www.genesdev.org/cgi/doi/10.1101/gad.270611.115

Erxleben A, Gessler A, Vervliet-Scheebaum M, Reski R (2012) Metabolite profiling of the moss *Physcomitrella patens* reveals evolutionary conservation of osmoprotective substances. Plant Cell Rep 31:427–436. 10.1007/s00299-011-1177-9

Fernandez-Pozo N, Haas FB, Meyberg R, Ullrich KK, Hiss M, Perroud P-F, Hanke S, Kratz V, Powell AF, Vesty EF et al. (2020) PEATmoss (Physcomitrella Expression Atlas Tool): a unified gene expression atlas for the model plant *Physcomitrella patens*. Plant J 102:165–177. 10.1111/tpj.14607

Fesenko I, Kirov I, Kniazev A, Khazigaleeva R, Lazarev V, Kharlampieva D, Grafskaia E, Zgoda V, Butenko I, Arapidi G et al. (2019) Distinct types of short open reading frames are translated in plant cells. Genome Res 29:1464–1477. 10.1101/gr.253302.119

Fesenko I, Shabalina SA, Mamaeva A, Knyazev A, Glushkevich A, Lyapina I, Ziganshin R, Kovalchuk S, Kharlampieva D, Lazarev V et al. (2021) A vast pool of lineage-specific microproteins encoded by long non-coding RNAs in plants. Nucleic Acids Res 49:10328–10346. 10.1093/nar/gkab816

Fortelny N, Yang S, Pavlidis P, Lange PF, Overall CM (2015) Proteome TopFIND 3.0 with TopFINDer and PathFINDer: database and analysis tools for the association of protein termini to pre-and post-translational events. Nucleic Acids Res 43:D290–D297. 10.1093/nar/gku1012

Fujino T, Kojima M, Beppu M, Kikugawa K, Yasuda H, Takahashi K (2000) Identification of the cleavage sites of oxidized protein that are susceptible to oxidized protein hydrolase (OPH) in the primary and tertiary structures of the protein. J Biochem 127:1087–1093. 10.1093/oxfordjournals.jbchem.a022702

Fuss J, Liegmann O, Krause K, Rensing SA (2013) Green targeting predictor and ambiguous targeting predictor 2: the pitfalls of plant protein targeting prediction and of transient protein expression in heterologous systems. New Phytol 200:1022–1033. 10.1111/nph.12433

Giglione C, Meinnel T (2021) Evolution-driven versatility of N terminal acetylation in photoautotrophs. Trends Plant Sci 26:375–391. 10.1016/j.tplants.2020.11.012

Gleason AC, Ghadge G, Chen J, Sonobe Y, Roos RP (2022a) Machine learning predicts translation initiation sites in neurologic diseases with nucleotide repeat expansions. PLOS ONE 17:e0256411. 10.1371/journal.pone.0256411

Gleason AC, Ghadge G, Sonobe Y, Roos RP (2022b) Kozak similarity score algorithm identifies alternative translation initiation codons implicated in cancers. Int J Mol Sci 23:10564. 10.3390/ijms231810564

Goodstein DM, Shu S, Howson R, Neupane R, Hayes RD, Fazo J, Mitros T, Dirks W, Hellsten U, Putnam N, Rokhsar DS (2012) Phytozome: a comparative platform for green plant genomics. Nucleic Acids Res 40:D1178–D1186. 10.1093/nar/gkr944

Grimm R, Grimm M, Eckerskorn C, Pohlmeyer K, Röhl T, Soll J (1997) Postimport methylation of the small subunit of ribulose-1, 5-bisphosphate carboxylase in chloroplasts. FEBS Lett 408:350–354. 10.1016/S0014-5793(97)00462-6

Healey AL, Piatkowski B, Lovell JT, Sreedasyam A, Carey SB, Mamidi S, Shu S, Plott C, Jenkins J, Lawrence T, Aguero B, Carrell AA, Nieto-Lugilde M, Talag J, Duffy A, Jawdy S, Carter KR, Boston L-B, Jones T, Jaramillo-Chico J, Harkess A, Barry K, Keymanesh K, Bauer D, Grimwood J, Gunter L, Schmutz J, Weston DJ, Shaw AJ (2023) Newly identified sex chromosomes in the *Sphagnum* (peat moss) genome alter carbon sequestration and ecosystem dynamics. Nat Plants 9:238–254. 10.1038/s41477-022-01333-5

Heintz D, Erxleben A, High AA, Wurtz V, Reski R, Van Dorsselaer A, Sarnighausen E (2006) Rapid alteration of the phosphoproteome in the moss *Physcomitrella patens* after cytokinin treatment. J Proteome Res 5:2283–2293. 10.1021/pr060152e

Hoernstein SNW, Mueller SJ, Fiedler K, Schuelke M, Vanselow JT, Schuessele C, Lang D, Nitschke R, Igloi GL, Schlosser A, Reski R (2016) Identification of targets and interaction partners of arginyl-tRNA protein transferase in the moss *Physcomitrella patens*. Mol Cell Proteomics 15:1808–1822. 10.1074/mcp.M115.057190

Hoernstein SNW, Fode B, Wiedemann G, Lang D, Niederkrüger H, Berg B, Schaaf A, Frischmuth T, Schlosser A, Decker EL, Reski R (2018) Host cell proteome of *Physcomitrella patens* harbors proteases and protease inhibitors under bioproduction conditions. J Proteome Res 17:3749–3760. 10.1021/acs.jproteome.8b00423

Hoernstein SNW, Özdemir B, van Gessel N, Miniera AA, Rogalla von Bieberstein B, Nilges L, Schweikert Farinha J, Komoll R, Glauz S, Weckerle T et al. (2023) A deeply conserved protease, acylamino acid-releasing enzyme (AARE), acts in ageing in Physcomitrella and Arabidopsis. Comm Biol 6:61. 10.1038/s42003-023-04428-7

Hohe A, Decker EL, Gorr G, Schween G, Reski R (2002) Tight control of growth and cell differentiation in photoautotrophically growing moss (*Physcomitrella patens*) bioreactor cultures. Plant Cell Rep 20:1135–1140. 10.1007/s00299-002-0463-y

Hohe A, Egener T, Lucht JM, Holtorf H, Reinhard C, Schween G, Reski R (2004) An improved and highly standardized transformation procedure allows efficient production of single and multiple targeted gene-knockouts in a moss, *Physcomitrella patens*. Curr Genet 44:339–347. 10.1007/s00294-003-0458-4

Horst NA, Katz A, Pereman I, Decker EL, Ohad N, Reski R (2016) A single homeobox gene triggers phase transition, embryogenesis and asexual reproduction. Nat Plants 2:15209. 10.1038/nplants.2015.209

Hu R, Li X, Hu Y, Zhang R, Lv Q, Zhang M, Sheng X, Zhao F, Chen Z, Ding Y et al. (2023) Adaptive evolution of the enigmatic *Takakia* now facing climate change in Tibet. Cell 186:3558–3576. 10.1016/j.cell.2023.07.003

Huesgen PF, Alami M, Lange PF, Foster LJ, Schröder WP, Overall CM, Green, BR (2013) Proteomic amino-termini profiling reveals targeting information for protein import into complex plastids. PLOS ONE 8:e74483. 10.1371/journal.pone.0074483

Jumper J, Evans R, Pritzel A, Green T, Figurnov M, Ronneberger O, Tunyasuvunakool K, Bates R, Žídek A, Potapenko A, et al. (2021) Highly accurate protein structure prediction with AlphaFold. Nature 596:583–589. 10.1038/s41586-021-03819-2

Katoh K, Standley DM (2013) MAFFT multiple sequence alignment software version 7: improvements in performance and usability. Mol Biol Evol 30:772–780. 10.1093/molbev/mst010

Keller A, Nesvizhskii AI, Kolker E, Aebersold R (2002) Empirical statistical model to estimate the accuracy of peptide identifications made by MS/MS and database search. Anal Chem 74:5383–5392. 10.1021/ac025747h

Kiessling J, Martin A, Gremillon L, Rensing SA, Nick P, Sarnighausen E, Reski R (2004) Dual targeting of plastid division protein FtsZ to chloroplasts and the cytoplasm. EMBO Rep 5:889–894. 10.1038/sj.embor.7400238

Kirbis A, Waller M, Ricca M, Bont Z, Neubauer A, Goffinet B, Szövényi P (2020) Transcriptional landscapes of divergent sporophyte development in two mosses, *Physcomitrium* (Physcomitrella) *patens* and *Funaria hygrometrica*. Front Plant Sci 11: 747. 10.3389/fpls.2020.00747

Kleifeld O, Doucet A, auf dem Keller U, Prudova A, Schilling O, Kainthan RK, Starr AE, Foster LJ, Kizhakkedathu JN, Overall CM (2010) Isotopic labeling of terminal amines in complex samples identifies protein N-termini and protease cleavage products. Nat Biotech 28:281–288. 10.1038/nbt.1611

Knosp S, Kriegshauser L, Tatsumi K, Malherbe L, Erhardt M, Wiedemann G, Bakan B, Kohchi T, Reski R, Renault H (2024) An ancient role for CYP73 monooxygenases in phenylpropanoid biosynthesis and embryophyte development. EMBO J: 10.1038/s44318-024-00181-7

Kozlov AM, Darriba D, Flouri T, Morel B, Stamatakis A (2019) RAxML-NG: a fast, scalable and user-friendly tool for maximum likelihood phylogenetic inference. Bioinform 35:4453–4455. 10.1093/bioinformatics/btz305

Kunze M, Berger J (2015) The similarity between N-terminal targeting signals for protein import into different organelles and its evolutionary relevance. Front Physiol 6:159761. 10.3389/fphys.2015.00259

Lang D, Ullrich KK, Murat F, Fuchs J, Jenkins J, Haas FB, Piednoel M, Gundlach H, Van Bel M, Meyberg R et al. (2018) The *Physcomitrella patens* chromosome-scale assembly reveals moss genome structure and evolution. Plant J 93:515–533. 10.1111/TPJ.13801

Linster E, Wirtz M (2018) N-terminal acetylation: an essential protein modification emerges as an important regulator of stress responses. J Ex Bot 69:4555–4568. 10.1093/jxb/ery241

Lueth VM, Reski R (2023) Mosses. Curr Biol 33:R1175–R1181. 10.1016/j.cub.2023.09.042

Maclean J, Koekemoer M, Olivier AJ, Stewart D, Hitzeroth II, Rademacher T, Fischer R, Williamson AL, Rybicki EP (2007) Optimization of human papillomavirus type 16 (HPV-16) L1 expression in plants: comparison of the suitability of different HPV-16 L1 gene variants and different cell-compartment localization. J Gen Virol 88, 1460–1469. 10.1099/vir.0.82718-0

Mao L, Kawaide H, Higuchi T, Chen M, Miyamoto K, Hirata Y, Kimura H, Miyazaki S, Teruya M, Fujiwara K, Tomita K, Yamane H, Hayashi KI, Nojiri H, Jia L, Qiu J, Ye C, Timko MP, Fan L, Okada K (2020) Genomic evidence for convergent evolution of gene clusters for momilactone biosynthesis in land plants. Proc Natl Acad Sci USA 117:12472–12480. 10.1073/pnas.1914373117

Marienfeld JR, Reski R, Friese C, Abel WO (1989) Isolation of nuclear, chloroplast and mitochondrial DNA from the moss *Physcomitrella patens*. Plant Sci 61:235–244. 10.1016/0168-9452(89)90230-6

McDonald L, Beynon RJ (2006) Positional proteomics: preparation of amino-terminal peptides as a strategy for proteome simplification and characterization. Nat Protoc 1:1790–1798. 10.1038/nprot.2006.317

Medina R, Johnson MG, Liu Y, Wickett NJ, Shaw AJ, Goffinet B (2019) Phylogenomic delineation of *Physcomitrium* (Bryophyta: Funariaceae) based on targeted sequencing of nuclear exons and their flanking regions rejects the retention of *Physcomitrella*, *Physcomitridium* and *Aphanorrhegma*. J Syst Evol 57:404–417. 10.1111/JSE.12516

Meinnel T, Giglione C (2008) Tools for analyzing and predicting N-terminal protein modifications. Proteomics 8:626–649. 10.1002/pmic.200700592

Meinnel T, Giglione C (2022) N-terminal modifications, the associated processing machinery, and their evolution in plastid-containing organisms. J Exp Bot 73:6013–6033. 10.1093/jxb/erac290

Mueller SJ, Lang D, Hoernstein SNW, Lang EG, Schuessele C, Schmidt A, Fluck M, Leisibach D, Niegl C, Zimmer AD, Schlosser A, Reski R (2014) Quantitative analysis of the mitochondrial and plastid proteomes of the moss *Physcomitrella patens* reveals protein macrocompartmentation and microcompartmentation. Plant Physiol 164:2081–2095. 10.1104/pp.114.235754

Nakai A, Yamauchi Y, Sumi S, Tanaka K (2012) Role of acylamino acid-releasing enzyme/oxidized protein hydrolase in sustaining homeostasis of the cytoplasmic antioxidative system. Planta 236:427–436. 10.1007/s00425-012-1614-1

Nelson D, Salamini F, Bartels D (1994) Abscisic acid promotes novel DNA-binding activity to a desiccation-related promoter of *Craterostigma plantagineum*. Plant J 5:451–458. 10.1046/j.1365-313X.1994.05040451.x

Nesvizhskii AI, Keller A, Kolker E, Aebersold R (2003) A statistical model for identifying proteins by tandem mass spectrometry. Anal Chem 75:4646–4658. 10.1021/ac0341261

Ouyang S, Zhu W, Hamilton J, Lin H, Campbell M, Childs K, Thibaud-Nissen F, Malek RL, Lee Y, Zheng L et al. (2007) The TIGR Rice Genome Annotation Resource: improvements and new features. Nucleic Acids Res 35:D883–D887. 10.1093/nar/gkl976

Perez-Riverol Y, Bai J, Bandla C, García-Seisdedos D, Hewapathirana S, Kamatchinathan S, Kundu DJ, Prakash A, Frericks-Zipper A, Eisenacher M et al. (2022) The PRIDE database resources in 2022: a hub for mass spectrometry-based proteomics evidences. Nucleic Acids Res 50:D543–D552. 10.1093/nar/gkab1038

Perroud P-F, Haas FB, Hiss M, Ullrich KK, Alboresi A, Amirebrahimi M, Barry K, Bassi R, Bonhomme S, Chen H et al. (2018) The *Physcomitrella patens* gene atlas project: large-scale RNA-seq based expression data. Plant J 95:168–182. 10.1111/tpj.13940

Petkowski JJ, Bonsignore LA, Tooley JG, Wilkey DW, Merchant ML, Macara IG, Schaner Tooley, CE (2013) NRMT2 is an N-terminal monomethylase that primes for its homologue NRMT1. Biochem J 456:453–462. 10.1042/BJ20131163

Prochnik, SE, Umen, J, Nedelcu, AM, Hallmann, A, Miller, SM, Nishii, I, Ferris, P, Kuo, A, Mitros, T, Fritz-Laylin, LK et al. (2010) Genomic analysis of organismal complexity in the multicellular green alga *Volvox carteri*. Science 329:223–226. 10.1126/science.1188800

Purwaha P, Silva LP, Hawke DH, WeinsteinJN, Lorenzi PL (2014) An artifact in LC-MS/MS measurement of glutamine and glutamic acid: in-source cyclization to pyroglutamic acid. Anal Chem 86:5633–5637. 10.1021/ac501451v

R Core Team (2024) R: A Language and environment for statistical computing. R Foundation for Statistical Computing. https://www.R-project.org

Ree R, Varland S, Arnesen T (2018) Spotlight on protein N-terminal acetylation. Exp Mol Med 50:1–13. 10.1038/s12276-018-0116-z

Renault H, Alber A, Horst NA, Basilio Lopes A, Fich EA, Kriegshauser L, Wiedemann G, Ullmann P, Herrgott L, Erhardt M et al. (2017) A phenol-enriched cuticle is ancestral to lignin evolution in land plants. Nat Comm 8: 14713. 10.1038/ncomms14713

Rensing SA, Lang D, Zimmer AD, Terry A, Salamov A, Shapiro H, Nishiyama T, Perroud PF, Lindquist EA, Kamisugi Y et al. (2008) The *Physcomitrella* genome reveals evolutionary insights into the conquest of land by plants. Science 319:64–69. 10.1126/science.1150646

Reski R, Abel WO (1985) Induction of budding on chloronemata and caulonemata of the moss, *Physcomitrella patens*, using isopentenyladenine. Planta 165:354–358. 10.1007/BF00392232

Rowland E, Kim J, Bhuiyan NH, van Wijk KJ (2015) The Arabidopsis chloroplast stromal N-terminome: complexities of amino-terminal protein maturation and stability. Plant Physiol 169:1881–1896. 10.1104/pp.15.01214

Ruiz-Molina N, Parsons J, Müller M, Hoernstein SNW, Bohlender LL, Pumple S, Zipfel PF, Häffner K, Reski R, Decker EL (2022) A synthetic protein as efficient multitarget regulator against complement over-activation. Comm Biol 5:152. 10.1038/s42003-022-03094-5

Sarnighausen E, Wurtz V, Heintz D, van Dorsselaer A, Reski R (2004) Mapping of the *Physcomitrella patens* proteome. Phytochem 65:1589–1607. 10.1016/j.phytochem.2004.04.028

Schaaf A, Tintelnot S, Baur A, Reski R, Gorr G, Decker EL (2005) Use of endogenous signal sequences for transient production and efficient secretion by moss (*Physcomitrella patens*) cells. BMC Biotech 5:30. 10.1186/1472-6750-5-30

Schaner Tooley CE, Petkowski JJ, Muratore-Schroeder TL, Balsbaugh JL, Shabanowitz J, Sabat M, Minor D, Hunt DF, Macara IG (2010) NRMT is an α-N-methyltransferase that methylates RCC1 and retinoblastoma protein. Nature 466:1125–1128. 10.1038/nature09343

Schilling S, Wasternack C, Demuth HU (2008) Glutaminyl cyclases from animals and plants: a case of functionally convergent protein evolution. Biol Chem 389:983–991. 10.1515/BC.2008.111

Schween G, Hohe A, Koprivova A, Reski R (2003) Effects of nutrients, cell density and culture techniques on protoplast regeneration and early protonema development in a moss, *Physcomitrella patens*. J Plant Physiol 160:209–212. 10.1078/0176-1617-00855

Shimizu K, Fujino T, Ando K, Hayakawa M, Yasuda H, Kikugawa K (2003) Overexpression of oxidized protein hydrolase protects COS-7 cells from oxidative stress-induced inhibition of cell growth and survival. Biochem Biophys Res Comm 304:766–771. 10.1016/S0006-291X(03)00657-0

Sperschneider J, Catanzariti AM, DeBoer K, Petre B, Gardiner DM, Singh KB, Dodds PN, Taylor, JM (2017) LOCALIZER: subcellular localization prediction of both plant and effector proteins in the plant cell. Sci Rep 7:44598. 10.1038/srep44598

Staes A, Impens F, Van Damme P, Ruttens B, Goethals M, Demol H, Timmerman E, Vandekerckhove J, Gevaert K (2011) Selecting protein N-terminal peptides by combined fractional diagonal chromatography. Nat Protoc 6:1130–1141. 10.1038/nprot.2011.355

Stock A, Clarke S, Clarke C, Stock J (1987) N-terminal methylation of proteins: structure, function and specificity. FEBS Lett 220:8–14. 10.1016/0014-5793(87)80866-9

Tasaki T, Sriram SM, Park KS, Kwon YT (2012) The N-end rule pathway. Annu Rev Biochem 81:261–289. 10.1146/annurev-biochem-051710-093308

Teixeira PF, Kmiec B, Branca RMM, Murcha MW, Byzia A, Ivanova A, Whelan J, Drag M, Lehtiö J, Glaser E (2017) A multi-step peptidolytic cascade for amino acid recovery in chloroplasts. Nat Chem Biol 13:15–17. 10.1038/nchembio.2227

Tschongov T, Konwar S, Busc, A, Sievert C, Hartmann A, Noris M, Gastoldi S, Aiello S, Schaaf A, Panse J, et al. (2024) Moss-produced human complement factor H with modified glycans has an extended half-life and improved biological activity. Front Immun 15:1383123. 10.3389/fimmu.2024.1383123

Tsunasawa S, Narita K, Ogata K (1975) Purification and properties of acylamino acid-releasing enzyme from rat liver. J Biochem 77:89–102. 10.1093/oxfordjournals.jbchem.a130722

van Kempen M, Kim SS, Tumescheit C, Mirdita M, Lee J, Gilchrist CL, Söding J, Steinegger M (2023) Fast and accurate protein structure search with Foldseek. Nat Biotech 42:243–246. 10.1038/s41587-023-01773-0

van Wijk KJ (2024) Intra-chloroplast proteases: A holistic network of chloroplast proteolysis. Plant Cell:koae178. 10.1093/plcell/koae178

Varadi M, Bertoni D, Magana P, Paramval U, Pidruchna I, Radhakrishnan M, Tsenkov M, Nair S, Mirdita M, Yeo J et al. (2024) AlphaFold Protein Structure Database in 2024: providing structure coverage for over 214 million protein sequences. Nucleic Acids Res 52:D368–D375. 10.1093/nar/gkad1011

Varshavsky A (1996) The N-end rule: functions, mysteries, uses. Proc Natl Acad Sci USA 93:12142–12149. 10.1073/pnas.93.22.12142

Varshavsky A (2019) N-degron and C-degron pathways of protein degradation. Proc Natl Acad Sci USA 116:358–366. 10.1073/pnas.181659611

Vuruputoor VS, Starovoitov A, Cai Y, Liu Y, Rahmatpour N, Hedderson TA, Wilding N, Wegrzyn JL, Goffinet B (2024) Crossroads of assembling a moss genome: navigating contaminants and horizontal gene transfer in the moss *Physcomitrellopsis africana*. G3 14:jkae104. 10.1093/g3journal/jkae104

Webb KJ, Lipson RS, Al-Hadid Q, Whitelegge JP, Clarke SG (2010) Identification of protein N-terminal methyltransferases in yeast and humans. Biochem 49:5225–5235. 10.1021/bi100428x

Wiedemann G, van Gessel N, Köchl F, Hunn L, Schulze K, Maloukh L, Nogué F, Decker EL, Hartung F, Reski R (2018) RecQ helicases function in development, DNA repair, and gene targeting in *Physcomitrella patens*. Plant Cell 30:717–736. 10.1105/tpc.17.00632

Wu R, Yue Y, Zheng X, Li H (2015) Molecular basis for histone N-terminal methylation by NRMT1. Gene Dev, 29:2337–2342. 10.1101/gad.270926.115

Yamauchi Y, Ejiri Y, Toyoda Y, Tanaka K (2003) Identification and biochemical characterization of plant acylamino acid–releasing enzyme. J Biochem 134:251–257. 10.1093/jb/mvg138

Yu G, Smith DK, Zhu H, Guan Y, Lam TTY (2017) ggtree: an R package for visualization and annotation of phylogenetic trees with their covariates and other associated data. Meth Ecol Evol 8:28–36. 10.1111/2041-210X.12628

Zhang J, Fu XX, Li RQ, Zhao X, Liu Y, Li MH, Zwaenepoel A, Ma H, Goffinet B, Guan YL, Xue JY, Liao YY, Wang QF, Wang QH, Wang JY, Zhang GQ, Wang ZW, Jia Y, Wang MZ, Dong SS, Yang J-F, Jiao YN, Guo YL, Kong HZ, Lu AM, Yang HM, Zhang SZ, Van de Peer Y, Liu ZJ, Chen ZD (2020) The hornwort genome and early land plant evolution. Nat Plants 6:107–118. 10.1038/s41477-019-0588-4

